# The REF6-dependent H3K27 demethylation establishes transcriptional competence to promote germination in *Arabidopsis*

**DOI:** 10.1101/2021.06.02.445236

**Authors:** Jie Pan, Huairen Zhang, Zhenping Zhan, Ting Zhao, Danhua Jiang

**Author notes:** Equal contribution (Jie Pan, Huairen Zhang).

## Abstract

Seed germination is a critical developmental switch from a dormant state to active growth, which involves extensive changes in metabolism, gene expression and cellular identity. However, our understanding of epigenetic and transcriptional reprogramming during this process is limited. The histone H3 lysine 27 trimethylation (H3K27me3) plays a key role in regulating gene repression and cell fate specification. Here, we profile H3K27me3 dynamics and dissect the function of H3K27 demethylation during germination. Our temporal genome-wide profiling of H3K27me3 and transcription reveal delayed H3K27me3 reprogramming compared with transcriptomic changes during germination, with H3K27me3 changes mainly occurring when the embryo is entering into vegetative development. REF6-mediated H3K27 demethylation promotes germination but does not significantly contribute to H3K27me3 dynamics during germination, but rather stably establishes an H3K27me3-depleted state permissive to transcription. By analyzing REF6 genomic binding, we show that it is absent from mature embryo chromatin and gradually establishes occupancy during the course of germination to counteract increased PRC2 activity. Our study provides key insights into the dynamics of gene expression and H3K27me3 during seed germination and suggests the function of H3K27me3 in facilitating cell fate switch. Furthermore, we reveal the importance of H3K27 demethylation-established transcriptional competence in germination and likely other developmental processes.

## Introduction

Histone H3 lysine 27 trimethylation (H3K27me3) is a repressive histone modification crucial for gene silencing and cell fate preservation (Wiles and Selker, 2017). In *Drosophila*, impaired maintenance of H3K27me3 leads to de-repression of *HOX* genes and failed cell lineage specification (Jurgens, 1985; Coleman and Struhl, 2017). In *Arabidopsis*, a large number of H3K27me3-marked genes are key for development and become specifically activated or repressed during the course of differentiation (Zhang et al., 2007; Lafos et al., 2011). Reprogramming of plant cell identity is coupled with re-distribution of H3K27me3 (Lee et al., 2019). A complete loss of H3K27me3 causes a loss of plant cell fate and induces somatic embryos and callus formation (Chanvivattana et al., 2004; Bouyer et al., 2011; Ikeuchi et al., 2015; Mozgova et al., 2017), suggesting an essential function for H3K27me3 in the establishment of plant cellular identity.

Plant H3K27me3 is deposited by the highly conserved Polycomb repressive complex 2 (PRC2) (Liu et al., 2010; Mozgova and Hennig, 2015), and is removed by JUMONJI (JMJ) DOMAIN-CONTAINING proteins, namely RELATIVE OF EARLY FLOWERING 6 (REF6), EARLY FLOWERING 6 (ELF6), JMJ13, JMJ30, and JMJ32 (Lu et al., 2011; Crevillen et al., 2014; Gan et al., 2014; Yan et al., 2018; Zander et al., 2019; Zheng et al., 2019). It is anticipated that these writers and erasers regulate H3K27me3 dynamics during development to facilitate cell fate specification. So far, they have been shown to regulate a series of developmental processes, such as the floral transition (Noh et al., 2004; Jiang et al., 2008; Gan et al., 2014; Hou et al., 2014; Li et al., 2018; Zheng et al., 2019), lateral root formation (Gu et al., 2014; Wang et al., 2019), leaf senescence (Wang et al., 2019), floral development (Katz et al., 2004; Liu et al., 2011; Sun et al., 2014; Yan et al., 2018), spermatogenesis (Borg et al., 2020), and embryogenesis (Grossniklaus et al., 1998).

The transition from seed dormancy to germination is one of the major events in plant development. In agricultural practice, quick and robust germination is usually a desired trait for synchronous growth after germination (Kermode, 2005). To ensure plants grow in their favorable seasons, germination is properly timed by both endogenous signals and environmental cues. One major type of endogenous signals is plant hormones. At the end of seed maturation, abscisic acid (ABA) is synthesized at high levels to induce seed desiccation and dormancy (Finkelstein et al., 2002). The biosynthesis and effects of ABA are antagonized by gibberellins (GA), which inhibit dormancy and promote germination (Nee et al., 2017). During germination, GA biosynthesis and ABA catabolic genes are activated, leading to increased GA and reduced ABA (Finkelstein et al., 2008). Accordingly, mutants defective in GA synthesis or ABA catabolism show reduced germination efficiency (Ogawa et al., 2003; Okamoto et al., 2006). Besides GA and ABA, additional hormones have been identified that regulate seed dormancy and germination. For instance, ethylene and brassinolide induce seed germination by counteracting the inhibitory effects of ABA (Steber and McCourt, 2001; Corbineau et al., 2014). Auxin, when applied at low or high concentrations, promotes or represses seed germination, respectively (Hsueh and Lou, 1947; Brady et al., 2003; Liu et al., 2007; He et al., 2012; Wang et al., 2016). Hormone-induced plant growth is linked with expansin, a family of proteins involved in cell wall loosening and extension (Cosgrove, 2015). During germination, the emerging radicle needs to break the restrictions imposed by its surrounding endosperm and testa (seed coat) (Holdsworth et al., 2008). Expansin promotes the weakening of these constraints, facilitating embryo growth and protrusion (Marowa et al., 2016).

Seed germination is coupled with major chromatin reconfiguration (van Zanten et al., 2011). H2B monoubiquitination (H2Bub1) activates the expression of a key dormancy gene *DELAY OF GERMINATION 1* (*DOG1*) (Liu et al., 2007). In contrast, *DOG1* expression is repressed by H3K9 methylation and H3K4 demethylation (Zheng et al., 2012; Zhao et al., 2015). Histone deacetylase 9 (HDA9) prevents germination through the repression of photosynthesis genes such as *RUBISCO SMALL SUBUNIT 2B* (*RBCS2B*) and *RUBISCO ACTIVASE* (*RCA*) (van Zanten et al., 2014). Moreover, loss of H3K27me3 in PRC2 subunit mutants causes enhanced dormancy and delayed germination (Bouyer et al., 2011). ALFIN1-like (AL) plant homeodomain (PHD)-containing proteins interact with the PRC1 complex and facilitate switching of chromatin state from permissive histone H3 Lysine 4 trimethylation (H3K4me3) to repressive H3K27me3 at seed-trait genes, including *DOG1, ABA INSENSITIVE 3* (*ABI3*), and *CHOTTO1* (*CHO1*), and thus repressing their expression to promote germination (Molitor et al., 2014). A recent study has also shown that REF6 activates the expression of ABA catabolic genes *CYP707A1* and *CYP707A3* in developing seeds by removing H3K27me3 at their loci, leading to reduced ABA levels and seed dormancy (Chen et al., 2020). However, the genome-wide dynamics of H3K27me3 and the role of active H3K27 demethylation during seed germination are still unclear.

Here, we show that REF6-mediated H3K27 demethylation promotes germination by facilitating transcriptional activation of hormone-related and expansin-coding genes during imbibition. Genome-wide H3K27me3 reprogramming only occurs when the embryo is switching to vegetative growth, despite large-scale transcriptomic changes at the initial stages of germination. Moreover, REF6 stimulates germination not by modulating H3K27me3 dynamics but rather by stably establishing an H3K27me3-depleted state to allow gene activation during germination. REF6 is absent from mature embryo chromatin, and its genomic binding is gradually established during germination, likely to prevent its targets from high levels of H3K27me3 mediated by the increased PRC2 activity. Our findings demonstrate a close link between H3K27me3 dynamics and cell fate transitions, and pinpoint a key role for REF6 in maintaining a constant H3K27me3-depleted environment for transcriptional competence.

## Results

### REF6 promotes seed germination

While screening for mutants with altered salt tolerance, we noted that after-ripened and stratified *ref6-1* seeds germinated slower than the wild-type (WT) Columbia (Col) after 3 days or 5 days on 1/2MS medium supplied with NaCl (Supplemental Fig. 1A and B). Similar phenotypes were observed with two other *ref6* mutant alleles *ref6-5* and *ref6*^*c*^ (a *ref6* null mutant generated by CRISPR-Cas9) (Yan et al., 2018) (Supplemental Fig. 1A and B). ELF6 and JMJ13 are close homologs of REF6. However, mutants defective in *ELF6, JMJ13*, or both displayed no strong germination defects under salt treatment (Supplemental Fig. 1A-C). Despite this, *REF6, ELF6* or *JMJ13* depletion had no impact on primary root growth under salt treatment relative to WT (Supplemental Fig. 1D). This suggested that REF6 might regulate germination independent of salt response.

We next closely examined germination of after-ripened seeds under normal conditions without salt treatment. Unlike *elf6* or *jmj13* mutants, all *ref6* alleles exhibited a delayed germination (Fig. 1A and B). Although *ref6* mutants germinated slower than WT, nearly 100% of seeds germinated after 4 days imbibition (Fig. 1B). Seed dormancy can be released by either extended storage (after-ripening) or imbibition at specific temperatures (stratification) (Nonogaki, 2014). To assess if delayed germination of *ref6* mutants is due to strong dormancy that cannot be fully released after long-term storage, after-ripened seeds were further stratified at 4°C for 3 days before subjected to germination, and similar phenotypes were observed with stratified seeds (Fig. 1C), suggesting that REF6 regulates germination independent of dormancy regulation. Nevertheless, we cannot exclude the possibility that dormancy induced by *REF6* depletion is immune to both after-ripening and stratification treatments. The delayed germination of *ref6-1* was largely complemented by HA-tagged REF6 (Fig. 1D) (Lu et al., 2011), confirming the role of REF6 in controlling germination. Since *ref6*^*c*^ is a null allele and all *ref6* mutants showed similar germination phenotypes, we used *ref6*^*c*^ for subsequent analyses.

**Figure 1.**
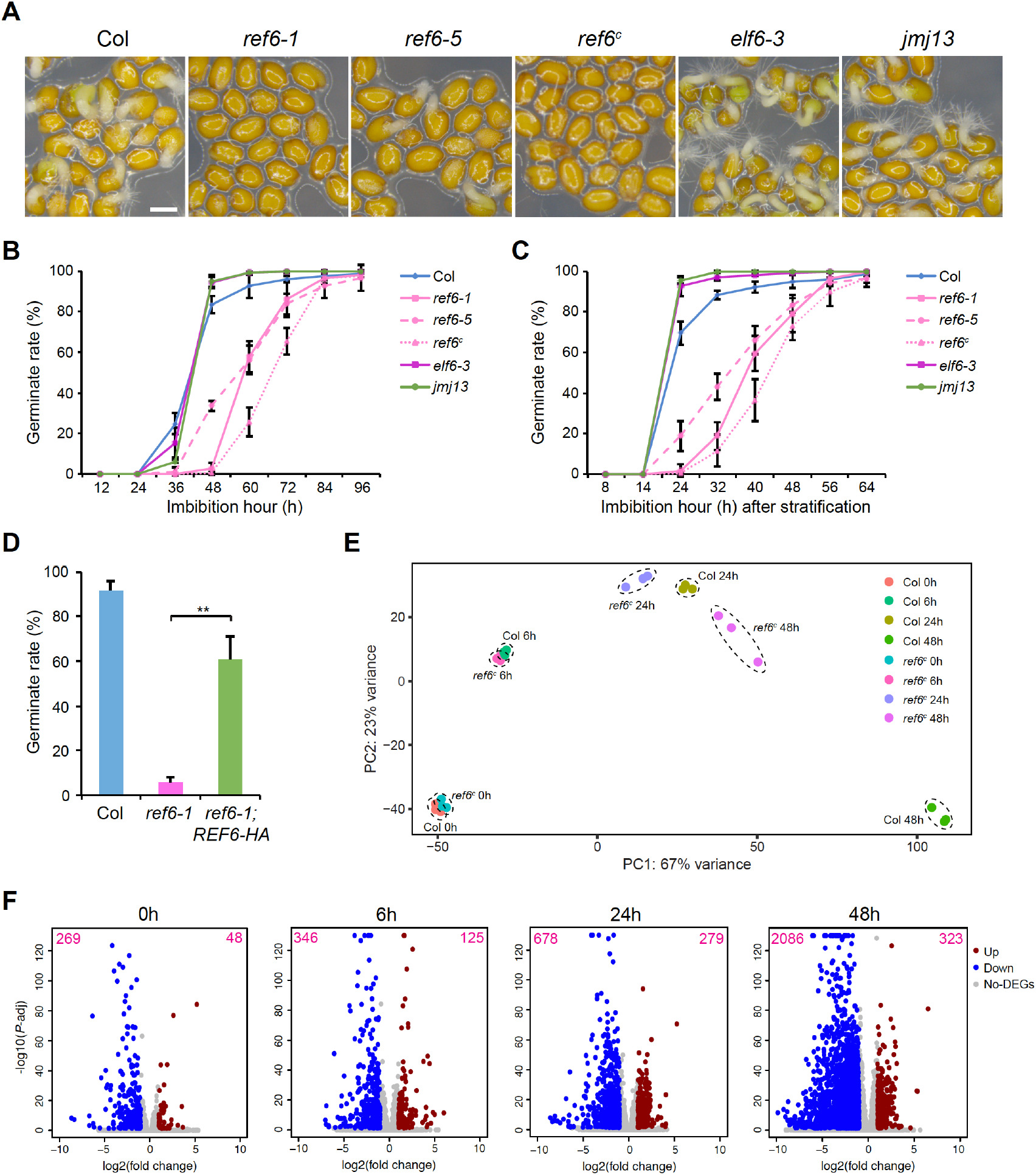
REF6 activates transcription to promote germination. **A**. The germination phenotype of after-ripened seeds imbibed for 48h on 1/2MS medium, Scale bar=0.5cm. **B, C**. The germination rates of after-ripened seeds (B) and after-ripened and stratified seeds (C). Values are means ± SD of three biological repeats. At least 50 seeds were analyzed for each replicate. **D** The germination rates of after-ripened seeds imbibed for 48h. Values are means ± SD of three biological repeats. At least 50 seeds were analyzed for each replicate. Statistical significance was determined by two-tailed Student’s *t*-test (**, *P*<0.01). **E** PCA plot of Col and *ref6*^c^ seeds imbibed for different times showing their transcriptome differences determined by RNA-seq. PC1 covers the highest amount of variance between samples; PC2 covers most of the remaining variance. Three biological replicates were performed for each line at each time point. **F** Volcano plots of differentially expressed genes (DEGs) for *ref6*^*c*^ and Col seeds imbibed for different times. The y-axis values correspond to −log_10_ (*P*-adjust), and the x-axis values correspond to log_2_ (fold change). Genes with at least two-fold expression changes and *P*-adjust less than 0.05 are considered as mis-expressed. The numbers of up- and down-regulated genes in *ref6*^*c*^ compared with Col were indicated at the top right and left corners respectively.

### REF6 activates transcription during germination

To evaluate the impact of REF6 on transcriptional regulation during germination, we performed RNA sequencing (RNA-seq) experiments with seeds imbibed for 0, 6, 24 and 48 hours (h). Principle component analysis (PCA) showed that the variation between *ref6*^*c*^ and WT transcriptomes was negligible at 0h but gradually increased as germination progressed (Fig. 1E). Diverging transcriptomes were also reflected by the numbers of differentially expressed genes (>2-fold changes) between *ref6*^*c*^ and WT, which increased from about 300 at 0h to over 2000 at 48h (Fig. 1F) (Supplemental Dataset 1). Most mis-expressed genes were down-regulated in *ref6*^*c*^ compared with WT (Fig. 1F), consistent with the role of REF6 in removing repressive H3K27me3 to activate gene expression. Together, these results demonstrate that REF6 is required to promote germination by activating gene expression.

### Large-scale H3K27me3 dynamics is coupled with cell fate switch during germination, following gene expression changes

Germination is a major developmental transition that involves extensive changes at physiological and molecular levels. Large-scale transcriptome reprogramming was indicated by drastic changes in gene expression, with 8417, 10453 and 11014 genes being differentially expressed in 6h, 24h and 48h imbibed WT seeds compared to the 0h control time point, respectively (Fig. 2A). We thus examined genome-wide dynamics of H3K27me3 using chromatin immunoprecipitation sequencing (ChIP-seq) to address its role in germination. Of the 8,949 H3K27me3 peaks recovered from 0h, 6h and 24h imbibed seeds, only 0.7% (67 peaks; 67 genes) and 0.6% (51 peaks; 42 genes) exhibited substantial changes (>2-fold) in the levels of H3K27me3 after 6h and 24h imbibition, respectively (Supplemental Fig. 2A) (Supplemental Dataset 2). In addition, most of these genes carried low levels of H3K27me3 compared with the average levels over all H3K27me3-marked genes (Supplemental Fig. 2B and C). Thus, despite strong changes in gene expression at 6h and 24h, the landscape of H3K27me3 was generally static at these time points.

**Figure 2.**
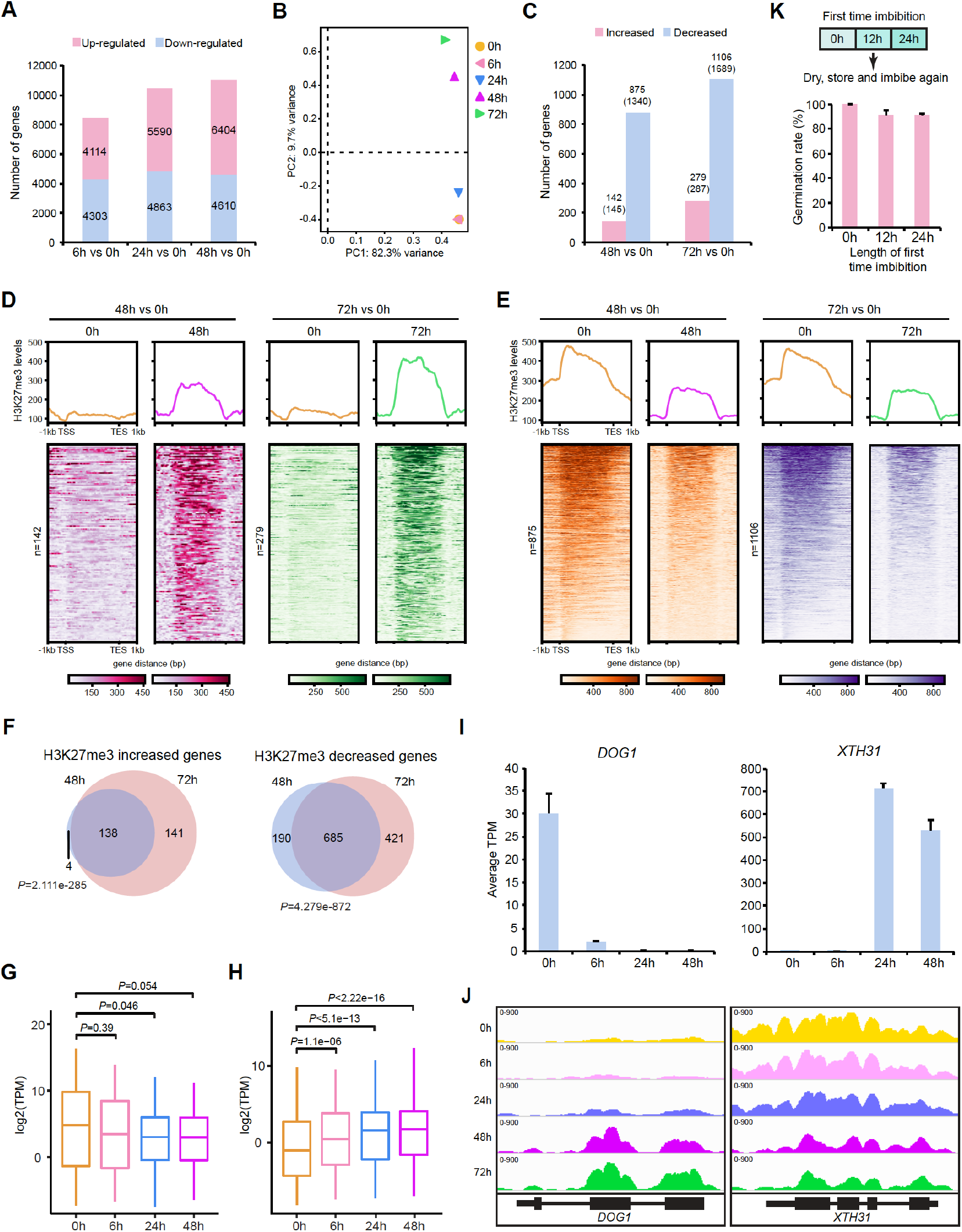
Dynamics of transcription and H3K27me3 during germination. **A** Numbers of up-regulated and down-regulated genes in imbibed Col seeds compared with 0h imbibed seeds determined by RNA-seq. **B** PCA plot of H3K27me3 profiles in Col seeds during germination determined by ChIP-seq. PC1 covers the highest amount of variance between samples; PC2 covers most of the remaining variance. The plot was generated after merging two biological replicates at each time point. **C** Numbers of H3K27me3 increased and decreased genes in 48h and 72h imbibed Col seeds compared with 0h imbibed seeds determined by ChIP-seq. Numbers in brackets show the amount of differential H3K27me3 peaks identified by ChIP-seq. **D** Average profiles and heat maps of normalized H3K27me3 ChIP-seq signals over H3K27me3 increased genes in 48h (n=142) and 72h (n=279) imbibed Col seeds compared with 0h imbibed seeds respectively. TSS: transcription start site, TES: transcription end site. The profiles were generated after merging two biological replicates at each time point. **E** Average profiles and heat maps of normalized H3K27me3 ChIP-seq signals over H3K27me3 decreased genes in 48h (n=875) and 72h (n=1106) imbibed Col seeds compared with 0h imbibed seeds respectively. The profiles were generated after merging two biological replicates at each time point. **F** Venn diagrams of H3K27me3 increased or decreased genes in 48h and 72h imbibed Col seeds compared with 0h imbibed seeds respectively. *P* values are calculated with hypergeometric test. **G, H**. Overall gene expression profiles of H3K27me3 increased (G) or decreased (H) genes identified in 48h imbibed Col seeds at different germination stages. The expression values represent average of three independent RNA-seq replicates. *P* values are based on the Mann-Whitney U test. **I** Expression of *DOG1* and *XTH31* in Col seeds imbibed for different times determined by RNA-seq. Values are means ± SD of three biological repeats. **J** Genome browser view of H3K27me3 enrichment at *DOG1* and *XTH31* loci in Col seeds imbibed for different hours. The profiles were generated after merging two biological replicates at each time point. **K** Germination rates of seeds imbibed once for different times. Imbibed seeds were collected, dried and stored for one week at room temperature, and subjected to germination once again. The rates of germination were measured 5 days after second time imbibition. Values are means ± SD of three biological repeats. At least 50 seeds were analyzed for each replicate.

We next assessed H3K27me3 changes at later germination phases by comparing H3K27me3 profiles from seeds imbibed for 48h and 72h (late phase) with those at 0h, 6h and 24h (early phase). Global H3K27me3 profiles in late phase seeds were relatively close and distinct from early phase seeds (Fig. 2B). Over 2000 peaks covering more than 1500 genes had differential H3K27me3 enrichment in late phase seeds, with the majority of loci showing reduced H3K27me3 (Fig. 2C-E) (Supplemental Dataset 2). H3K27me3 levels at these genes remained unchanged at 6h, while small changes were observed at 24h, particularly at genes where H3K27me3 was reduced (Supplemental Fig. 3). Genes with differential H3K27me3 identified at 48h and 72h significantly overlapped (Fig. 2F), suggesting that the major H3K27me3 changes initiated between 24h to 48h were maintained. During germination, seed-traits like dormancy are repressed while seedling-traits like biosynthesis are activated. Consistent with this developmental switch, loci that gained H3K27me3 were enriched for seed maturation and dormancy genes, while those with decreased H3K27me3 were mainly involved in transcription, biosynthesis and plant development (Supplemental Fig. 4).

To evaluate the impact of H3K27me3 changes on gene expression, we focused on genes with differential H3K27me3 enrichment at 48h and examined their expression during germination. The overall expression changes of these genes correlated with the H3K27me3 dynamics at their loci (Fig. 2G and H). Interestingly, we found that their transcripts levels decreased or increased prior to detectable H3K27me3 changes at 48h (Fig. 2G and H). For instance, although *DOG1* expression was repressed from 6h, H3K27me3 accumulated at the *DOG1* locus only after 48h of imbibition (Fig. 2I and J). Similarly, activation of a xyloglucan endo-transglycosylase coding gene *XTH31* started from 24h, prior to any reduction in H3K27me3 at this locus (Fig. 2I and J).

Notably, WT seeds start to germinate after imbibed for 24 to 48 hours (Fig. 1B), coinciding with the large-scale H3K27me3 changes. Because H3K27me3 is closely linked with cell fate specification, we reasoned that the H3K27me3 dynamics could be associated with the embryonic to vegetative cell fate switch, while gene expression changes are not. Seeds imbibed until 24h were collected, dried and stored for one week prior to sowing and germination once again (Fig. 2K). 24h imbibed seeds germinated normally after re-desiccation and storage (Fig. 2K), suggesting that the imbibed embryo is able to switch back to a dormant state at this time point, despite strong transcriptomic changes during imbibition. Hence, it is likely that during germination, H3K27me3 reprogramming only occurs when the embryo is committed to exiting dormancy and entering seedling development, after gene expression changes.

### REF6 demethylation establishes an H3K27me3-depleted state to promote gene activation during germination

Since the majority of H3K27me3 changes in late phase germinating seeds involved a reduction in H3K27me3, we further investigated the role of REF6 in regulating H3K27me3 dynamics during germination. Most WT but not *ref6*^*c*^ seeds displayed radicle protrusion after 48h imbibition (Fig. 1A and B). We thus excluded this time point because the variation in H3K27me3 between *ref6*^*c*^ and WT may be due to their different germination states. Genes that displayed reduced H3K27me3 at 72h in WT were examined for their H3K27me3 changes in 0h and 72h imbibed *ref6*^*c*^ seeds. However, loss of *REF6* did not significantly affect the depletion of H3K27me3 at these loci (Supplemental Fig. 5A-B), suggesting that REF6 is not essential for the H3K27me3 dynamics during germination.

To understand the function of REF6 on germination, we compared H3K27me3 profiles in *ref6*^*c*^ and WT. In 0h, 6h, and 24h imbibed seeds, 1529, 1818, and 1538 genes were hypermethylated (>2-fold increase) in *ref6*^*c*^ compared with WT, whereas only 14, 131 and 18 genes were hypomethylated (>2-fold decrease) respectively (Fig. 3A) (Supplemental Dataset 3), confirming the demethylation activity of REF6. Moreover, these hypermethylated genes significantly overlapped (Fig. 3B). We hereonin refer to all the hypermethylated genes from these three time points as “seed hypermethylated genes”. At every time point, these genes showed a clear increase of H3K27me3 in *ref6*^*c*^ (Fig. 3C and D). Thus, during the early phase of germination, REF6 is required for the H3K27me3-depleted state at a similar set of genes.

**Figure 3.**
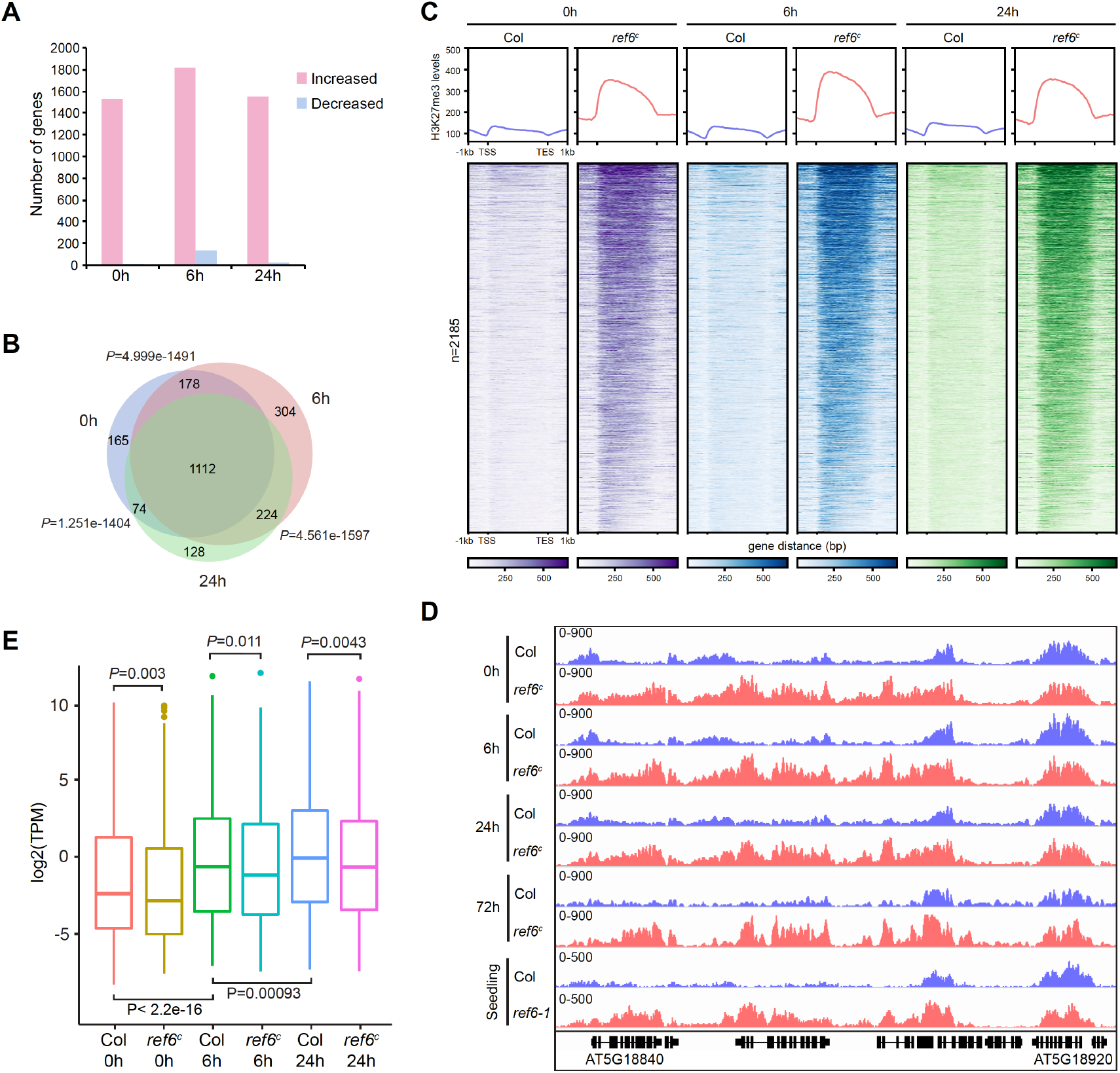
REF6 constantly mediates H3K27 demethylation at a similar set of genes to establish their transcriptional competence. **A** Numbers of H3K27me3 increased and decreased genes in 0h, 6h and 24h imbibed *ref6*^*c*^ seeds compared with same time imbibed Col seeds determined by ChIP-seq. **B** Venn diagram of H3K27me3 hypermethylated genes in 0h, 6h and 24h imbibed *ref6*^*c*^ seeds compared with same time imbibed Col seeds. *P* values are based on the hypergeometric test. **C** Average profiles and heat maps of normalized H3K27me3 ChIP-seq signals over a union set of seed hypermethylated genes (n=2185) in 0h, 6h and 24h imbibed Col and *ref6*^*c*^ seeds. The profiles were generated after merging two biological replicates at each time point. **D** Genome browser view of H3K27me3 enrichment in seeds imbibed for different hours at a genomic region showed previously with increased H3K27me3 levels in *ref6-1* seedlings (Cui et al., 2016). The profiles were generated after merging two biological replicates at each time point. **E** Overall gene expression profiles of seed hypermethylated genes at different germination stages in Col and *ref6*^*c*^. The expression values represent average of three independent RNA-seq replicates. *P* values are based on the Mann-Whitney U test.

To further examine the dynamics of REF6-mediated demethylation at later developmental stages, we compared hypermethylated genes in 0h imbibed *ref6*^*c*^ seeds with those in 72h imbibed *ref6*^*c*^ (n=1713) (Supplemental Dataset 3) and 12-day-old *ref6-1* seedlings (n=1688) identified from a previous study (Cui et al., 2016). These genes significantly overlapped (Supplemental Fig. 5C), while H3K27me3 enrichment profiles of the 0h, 72h and seedling-combined hypermethylated gene set showed a consistent increase of H3K27me3 in *ref6* mutants across datasets (Fig. 3D and Supplemental Fig. 5D). These results suggest that REF6 demethylated loci are largely shared from seeds to seedlings. In addition, among the 1106 genes losing H3K27me3 after 72h imbibition in WT, only 43 of them carried higher H3K27me3 levels in *ref6*^*c*^ compared with WT (Supplemental Fig. 5E), confirming that H3K27me3 reprogramming during germination is largely REF6 independent. Importantly, hypermethylated genes significantly overlapped with expression down-regulated genes in *ref6*^*c*^ seeds during germination (Supplemental Fig. 6). Seed hypermethylated genes tended to be activated during germination but the activation was compromised in *ref6*^*c*^, presumably due to increased H3K27me3 at these loci (Fig. 3E). Together, our results reveal stable REF6-dependent H3K27 demethylation during germination, which serves to derepress genes for activation.

### REF6 genomic binding is gradually established during germination

Genomic targeting of REF6 is mediated by its zinc-finger (ZnF) domain, which preferentially binds to unmethylated CTCTGYTY motifs (Y represents C or T) (Cui et al., 2016; Li et al., 2016; Qiu et al., 2019) at nearly 3000 genes in 12-day-old seedlings (Cui et al., 2016). To identify REF6 targets during germination, we first performed a pilot experiment to profile REF6 binding sites in 10-day-old seedlings using the *ref6-1;REF6-HA* transgenic line (Lu et al., 2011). 2272 peaks covering 2325 genes were bound by REF6 (Supplemental Dataset 4), which significantly overlapped with REF6 target genes previously identified in 12-day-old seedlings (Supplemental Fig. 7A) (Cui et al., 2016). Moreover, motif analysis at REF6 peaks obtained in our experiment successfully identified consensus sequences similar to CTCTGYTY (Supplemental Fig. 7B). Thus, our approach is sufficient to profile REF6 genomic targeting.

We next examined REF6 occupancy within seeds imbibed for different time periods. Surprisingly, only a few loci were bound by REF6 in 0h and 6h imbibed seeds, with the number of REF6 binding sites gradually increasing thereafter (Fig. 4A) (Supplemental Dataset 4). The majority of the REF6 target genes identified in 6h, 24h and 48h imbibed seeds overlapped with those in 10-day-old seedlings (Fig. 4B). Motif identification analysis discovered similar CTCTGYTY-like motifs bound by REF6, with the significance of enrichment increasing during germination (Fig. 4C). Consistently, almost all identified REF6 target genes in seeds contain the CTCTGYTY motif (Supplemental Fig. 7C). In addition, the accumulation of REF6 at its targets was enhanced from seeds to seedlings (Fig. 4D and E). These results suggest that REF6 genomic binding is gradually established during the course of germination. REF6 target genes identified in germinating seeds are enriched with seed hypermethylated genes in *ref6*^*c*^ (Supplemental Fig. 7D). However, REF6 target genes at 48h showed little overlap with genes losing H3K27me3 after 48h imbibition in WT (Supplemental Fig. 7E), further confirming that REF6 does not directly regulate H3K27me3 reprogramming during germination.

**Figure 4.**
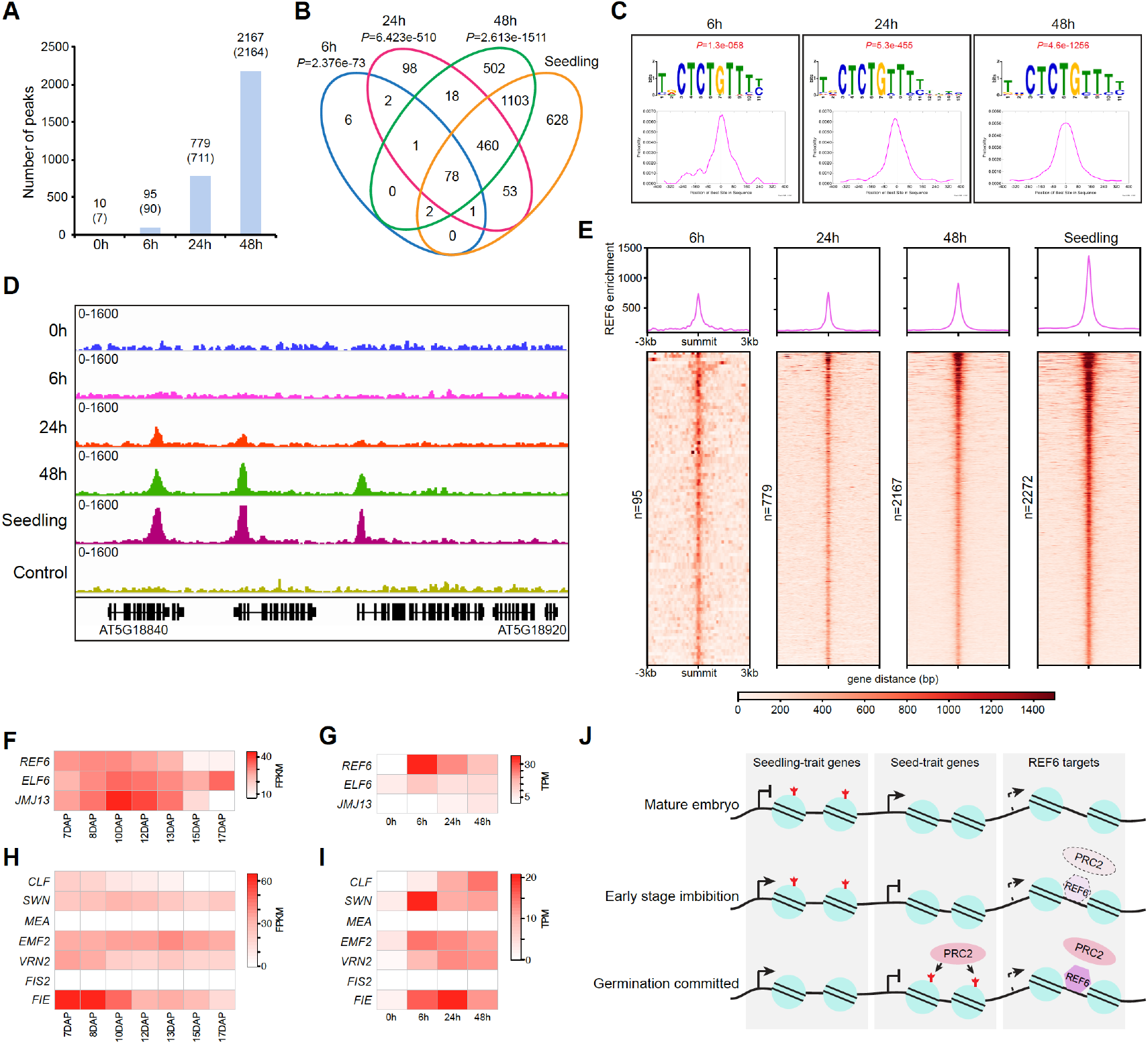
Establishment of REF6 genomic binding during germination. **A** Numbers of REF6 binding peaks during germination identified by ChIP-seq. Numbers in brackets show the amount of REF6 target genes identified by ChIP-seq. **B** Venn diagram of REF6 target genes in 6h, 24h, and 48h imbibed seeds and 10-day-old seedlings. *P* values are based on the hypergeometric test and are calculated to test the overlap significance between REF6 target genes in imbibed seeds at each time point and those in 10-day-old seedlings. **C** The top DNA motifs enriched and their distributions in REF6 binding peaks during germination. **D** Genome browser view of REF6-HA enrichment in seeds imbibed for different hours and 10-day-old seedlings at a genomic region showed in Fig. 3D. Col seedlings were used as a negative control for the HA ChIP. The profiles were generated after merging two biological replicates at each time point except for the negative control, which was performed with one replicate. **E** Average profiles and heat maps of normalized REF6-HA ChIP-seq signals around REF6 binding peak summits in 6h, 24h, and 48h imbibed seeds and 10-day-old seedlings. The profiles were generated after merging two biological replicates. **F, G**. Heat map showing expression levels of *REF6, ELF6*, and *JMJ13* during embryogenesis (F) and germination (G) in Col determined by RNA-seq. The expression levels represent average of three biological replicates. **H, I**. Heat map showing expression levels of PRC2 subunits during embryogenesis (H) and germination (I) in Col determined by RNA-seq. The expression levels represent average of three biological replicates. **J**. A possible model for the functions of transcription dynamics, the H3K27me3 reprogramming and REF6-mediated H3K27 demethylation during germination. In mature embryo, H3K27me3 represses the expression of seedling-trait genes, while seed-trait genes are highly expressed. At early phase germination, seedling-trait genes start to be activated and seed-trait genes are repressed, preparing seeds for germination. However, H3K27me3 landscape is preserved to safeguard the embryo identify at this stage. After long-term imbibition and when embryo is committed to entering vegetative development, H3K27me3 is removed from seedling-trait genes and meanwhile deposited at seed-trait genes to reinforce their silencing. REF6 and PRC2 subunits share similar expression patterns during embryogenesis and germination. Thus, although REF6 is absent from the mature embryo chromatin, low H3K27me3 levels and the transcriptional competence (indicated with dashed arrow) established by REF6 are maintained likely due to the lack of PRC2. During germination, the REF6 genomic binding is gradually established to counteract the increased PRC2 activity.

The lack of substantial REF6 binding within the chromatin of mature seeds led us to examine the expression of *REF6* during seed maturation and germination. *REF6* expression was reduced during embryo maturation and reactivated after 6h of imbibition (Fig. 4F and G), coinciding with the temporal initiation of REF6 binding across the genome. Notably, despite little to no REF6 binding, low H3K27me3 levels were maintained at thousands of REF6 target genes in early phase imbibed seeds (Fig. 3C). We reasoned that PRC2 activity might be low during these stages, preventing these genes from accumulating H3K27me3. Indeed, several PRC2 subunits shared similar expression patterns with REF6 (Fig. 4H and I). Thus, it is likely that the H3K27me3-depleted state is established before seed maturation, and stably maintained in dry and early phase imbibed seeds, where little PRC2 is present. When the embryo is committed to germination after a sufficient period of imbibition, REF6 occupancy at its targets is fully established to inhibit the high accumulation of H3K27me3 once PRC2 levels are restored (Fig. 4J).

### REF6 demethylates H3K27me3 at hormone-related and expansin-coding genes to promote germination

We next investigated the mechanism of how REF6 functions in stimulating seed germination. GA and ABA are two major hormones that antagonistically regulate germination, during which both GA synthesis and ABA catabolism are activated, resulting in increased GA and reduced ABA levels. Two GA synthetic genes (*GA20ox1* / 2) and two ABA catabolic genes (*CYP707A1* / 3) were targeted by REF6 binding and showed higher levels of H3K27me3 in *ref6*^*c*^ (Fig. 5A and Supplemental Fig. 8A), suggesting that they are direct REF6 targets. H3K27me3 removal by REF6 at these loci was also evident in seedlings (Fig. 5A and Supplemental Fig. 8A), indicating that these genes are ubiquitous targets of REF6. RNA-seq and RT-qPCR results revealed that the expression of *GA20ox1* and *CYP707A3*, but not *GA20ox2* and *CYP707A1*, was reduced in *ref6*^*c*^ seeds during germination (Fig. 5B and Supplemental Fig. 8B). Thus, REF6-mediated H3K27me3 removal at *GA20ox2* and *CYP707A1* is not required for their expression during germination, but may rather serve to prepare their activation at later developmental stages. Indeed, previous findings show that *GA20ox2* and *CYP707A1* are less expressed in *ref6* mutants at the seedling stage (Lu et al., 2011; Li et al., 2016).

**Figure 5.**
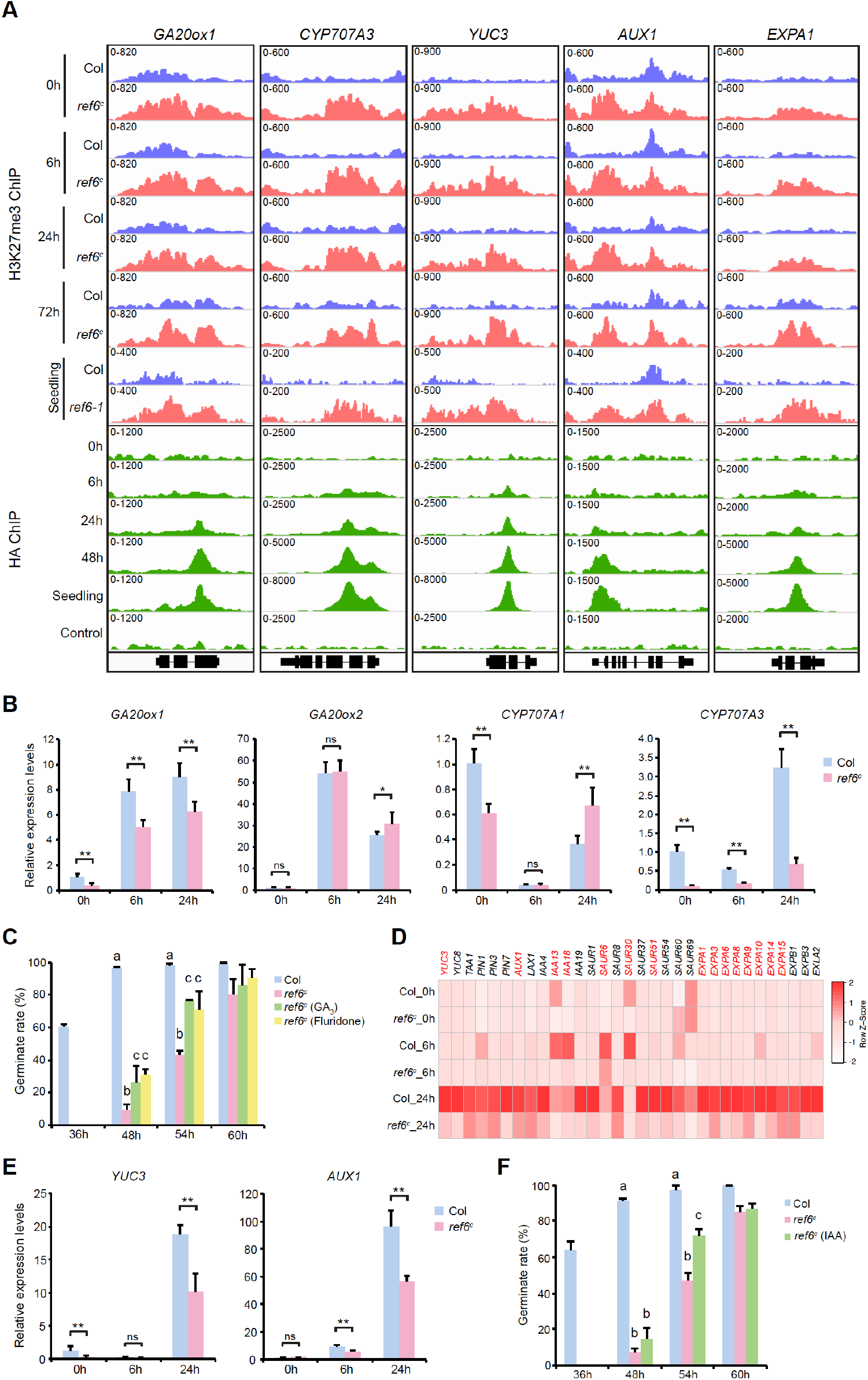
REF6 demethylates hormone-related and expansion coding genes to promote their expression during germination. **A** Genome browser views of H3K27me3 and REF6-HA enrichment at *GA20ox1, CYP7070A3, YUC3, AUX1*, and *EXPA1* loci in imbibed seeds and seedlings. Col seedlings were used as a negative control for the HA ChIP. The profiles were generated after merging two biological replicates at each time point except for the negative control, which was performed with one replicate. **B** Expression of *GA20ox1, GA20ox2, CYP707A1* and *CYP707A3* in Col and *ref6*^*c*^ during germination determined by RT-qPCR. Values are means ± SD of three biological repeats. Statistical significance was determined by two-tailed Student’s *t*-test (ns, not significant; *, *P*<0.05; **, *P*<0.01). **C** The germination rates of after-ripened seeds imbibed on 1/2MS medium with or without GA_3_ (10µM) and fluridone (2µM). Values are means ± SD of three biological repeats. At least 50 seeds were analyzed for each replicate. The significance of differences at 48h and 54h was tested using one-way ANOVA with Tukey’s test (*P*<0.05), different letters indicate statistical significant. **D** Heat map showing the relative expression changes (z-normalized) of auxin-related and expansin genes during germination determined by RNA-seq. The values used for generating heat map are average TPM of three independent RNA-seq replicates. Genes in red are REF6 direct targets identified by ChIP-seq. **E** Expression of *YUC3* and *AUX1* in Col and *ref6*^*c*^ during germination determined by RT-qPCR. Values are means ± SD of three biological repeats. Statistical significance was determined by two-tailed Student’s *t*-test (ns, not significant; **, *P*<0.01). **F** The germination rates of after-ripened seeds imbibed on 1/2MS medium with or without IAA (1nM). Values are means ± SD of three biological repeats. At least 50 seeds were analyzed for each replicate. The significance of differences at 48h and 54h was tested using one-way ANOVA with Tukey’s test (*P*<0.05), different letters indicate statistical significant.

We hypothesized that reduced expression of *GA20ox1* and *CYP707A3* might compromise GA synthesis and ABA catabolism during germination and thus lead to delayed germination of *ref6* seeds. We thus germinated fully after-ripened *ref6*^*c*^ seeds in the presence of GA_3_ and the ABA inhibitor fluridone. Both GA_3_ and fluridone stimulated the germination of *ref6*^*c*^ seeds (Fig. 5C), albeit at lower germination rates than that of WT. These results confirmed that REF6 promotes germination, at least partially, through the regulation of GA synthesis and ABA catabolism.

In addition to GA and ABA pathways, we noted that several genes associated with auxin synthesis, transport and signaling were also down-regulated in *ref6*^*c*^ seeds during germination (Fig. 5D). Among them, *YUC3, AUX1* and several others were bound by REF6 and were hypermethylated in *ref6*^*c*^ at both seed and seedling stages (Fig. 5A and Supplemental Fig. 8A) (Supplemental Dataset 3 and 4) (Cui et al., 2016). YUC3 is one of the flavin monooxygenase-like enzymes involved in auxin biosynthesis, while AUX1 is an auxin influx transporter. Mutation of *AUX1* causes weakly delayed germination, while *AUX1* overexpression accelerates germination (Wang et al., 2016), suggesting a positive role of AUX1 on seed germination. The reduced expression of *YUC3* and *AUX1* in *ref6*^*c*^ was further validated by RT-qPCR (Fig. 5E). Consistently, exogenous application of IAA partially complemented the delayed germination of *ref6*^*c*^ seeds (Fig. 5F). Hence, auxin deficiency also contributes to germination defects upon loss of *REF6*.

Expansin-induced cell wall loosening promotes radicle protrusion and germination. We found that 11 out of 36 expansin coding genes were less transcribed during germination in *ref6*^*c*^, among which 8 α-expansin (EXPA) genes were targeted for H3K27me3 removal by REF6 (Fig. 5A and D) (Supplemental Dataset 3 and 4). Most of these expansin-coding genes were strongly transcribed after 24h of imbibition, although their H3K27me3-depleted state was already established by REF6 in mature embryo, further suggesting the function of REF6 demethylation in establishing transcriptional competence. Taken together, these results indicate that REF6 protects hormone-related and expansin-coding genes from H3K27me3 accumulation to ensure timely germination.

## Discussion

Unlike animals, plants can pause a progression through their life cycle by entering into seed dormancy, often over several years. The decision to germinate and resume growth is thus crucial for plant survival. In this study, we reveal the dynamics and uncoupling of transcription and H3K27me3 during seed germination. Extensive transcriptomic changes occur soon after seeds are exposed to favorable germination conditions, while H3K27me3 reprogramming lags behind until the embryo commits to germination. Because transcriptional changes could be more flexible before H3K27me3 silencing is re-established, this may allow seeds to prepare for germination with a possibility to reverse back to a dormant state. After long-term imbibition and the ideal conditions for plant growth are perceived, the embryo reprograms H3K27me3 silencing to switch cell fate and allow the next phase of development to proceed (Fig. 4J). During germination, many seed-trait genes gain H3K27me3 (Supplemental Fig. 4A and C), these findings are in line with previous observations that PRC2 is vital for the embryonic to seedling phase switch (Bouyer et al., 2011).

The behavior of H3K27me3 dynamics during germination suggests that it is not closely linked with the initiation of gene repression, but is rather employed to reinforce a transcriptionally silenced state. Such a phenomenon is also observed during vernalization, when prolonged cold exposure induces *FLC* repression and localized H3K27me3 deposition around the transcription start site of *FLC*. Once plants return to warmer conditions suitable for flowering, H3K27me3 spreads across the whole *FLC* locus, resulting in stable *FLC* silencing and a commitment to flowering (Kim et al., 2009; Hepworth and Dean, 2015; Luo and He, 2020). Thus, our findings extend the H3K27me3 dynamics described at *FLC* to other loci, and indicate a general role of H3K27me3 in maintaining but not inducing transcriptional repression. Using mammalian cell lines, it has been shown that transcriptional inhibition occurs prior to H3K27me3 accumulation, while a lack of transcription is sufficient to induce H3K27me3 deposition (Riising et al., 2014; Hosogane et al., 2016). Gene repression may configure chromatin to a PRC2-friendly state, which enhances PRC2 binding and/or activity, resulting in high H3K27me3 levels that license the maintenance of gene silencing.

Our results reveal that a large number of seedling-trait genes lose H3K27me3 during germination. Similar H3K27me3 depletion observed in *ref6*^*c*^ (Supplemental Fig. 5A, B and E) and the lack of direct REF6 binding (Supplemental Fig. 7E) suggests that REF6 does not play a major role in removing H3K27me3 at these loci. It is unlikely that ELF6 and JMJ13 mediate this process since their mutation did not impact germination. The expression of these genes occurs before a strong reduction of H3K27me3, indicating the involvement of transcription factors in opposing the repressive effect of H3K27me3. In floral stem cells, homeotic transcription factor AGAMOUS (AG) competes with PRC2 for the same Polycomb response element (PRE) at their target gene *KNUCKLES* (*KNU*). AG binding causes the eviction of PRC2 and failed maintenance of H3K27me3 during cell division, facilitating the depletion of H3K27me3 at the *KNU* locus (Sun et al., 2014). Similarly, PRE excision in *Drosophila* embryo induces dilution of H3K27me3 in a cell cycle-dependent manner (Coleman and Struhl, 2017). At seedling-trait genes, H3K27me3 levels start to decrease between 24-48h after imbibition, coinciding with the activation of the cell cycle during germination (Supplemental Fig. 9) (Barroco et al., 2005). We thus hypothesize that H3K27me3 is passively diluted through cell divisions due to insufficient maintenance by PRC2, which is exploited by transcription factors to induce active chromatin environment changes. Alternatively, other H3K27 demethylases, such as JMJ30 and JMJ32, may be involved in removing H3K27me3 during germination.

Although H3K27 demethylases are anticipated to regulate H3K27me3 dynamics, REF6 appears to stably maintain an H3K27me3-depleted state at a similar set of genes from seeds to seedlings. We show that though not regulating H3K27me3 dynamics, the REF6-dependent H3K27me3 removal is required for gene activation during germination (Fig. 3E). REF6 contains a ZnF domain capable of recognizing the CTCTGYTY motif (Cui et al., 2016; Li et al., 2016). Hence, it could be stably recruited by DNA sequences and demethylates similar targets, where REF6 establishes a constant H3K27me3-depleted state to maintain their transcriptional competence during plant development and environmental responses. Nevertheless, several studies have shown transcription factor-mediated genomic targeting of REF6 (Yu et al., 2008; Smaczniak et al., 2012; Hou et al., 2014; Hyun et al., 2016). In these cases, REF6 may be recruited to modulate H3K27me3 dynamics in a transcription factor-dependent manner.

In this study, we have demonstrated that the activation of *GA20ox1, CYP707A3*, and several auxin-related and expansin genes during germination are directly regulated by REF6. Exogenous application of GA, auxin or ABA synthesis inhibitor fluridone could all partially rescue the delayed germination of *ref6*^*c*^, suggesting that REF6 impacts on GA, ABA and auxin pathways to regulate radicle protrusion and promote seed germination. It has recently reported that in developing seeds, REF6 binds to the *CYP707A1* and *CYP707A3* loci and mediates their H3K27me3 removal and transcription activation, limiting seed dormancy (Chen et al., 2020). Collectively, these results suggest that REF6 functions in both seed maturation and germination stages to reduce dormancy and promote germination.

At the end of seed maturation, the embryo becomes inactive and preserves energy until suitable environmental conditions for seedling development arise. Consistently, chromatin becomes compacted and transcriptional activity in mature seeds is reduced (Comai and Harada, 1990; van Zanten et al., 2011). Our results show that the expression of REF6 and several PRC2 components, including the essential and non-redundant subunit FIE, is reduced during embryo maturation, while REF6 is absent from the chromatin of the mature embryo. Despite this, the H3K27me3-depleted state regulated by REF6 is still maintained, suggesting that compacted chromatin and reduced PRC2 activity in dry seeds helps to prevent H3K27me3 from accumulating at REF6 targets. Alternatively, the ELF6 expressed in mature embryo maintains the low H3K27me3 levels at REF6 targets (Fig. 4F and G). However, the unaffected germination rate of *elf6* mutant seeds indicates that ELF6 may not play a major role in preventing H3K27me3 accumulation at REF6 targets. Germination induces REF6 transcription, leading to its progressive genomic binding during seed imbibition, which coincides with the accumulation of PRC2 subunits and chromatin decondensation (van Zanten et al., 2011). Thus, REF6 binds to chromatin once again to protect its targets from gaining H3K27me3 upon re-activation of PRC2 during germination (Fig. 4J). We propose that the lack of PRC2 and REF6, together with chromatin compaction, preserves the H3K27me3 landscape to a dormant state in dry seeds.

## Methods

### Plant materials and growth conditions

All *Arabidopsis* materials are in Col background, *ref6-1* (Lu et al., 2011), *ref6-5* (*Antunez-Sanchez et al*., *2020*), *ref6*^*c*^ (Yan et al., 2018), *elf6-3* (Yu et al., 2008), *jmj13* (Zheng et al., 2019) and *ref6-1;REF6-HA* (Lu et al., 2011) were described previously. Plants were grown in long days (16h light/8h dark) at ~22°C.

### Seed germination test

After-ripened seeds (seeds harvested and stored at room temperature for at least three months) collected at the same time and growth conditions were sown on 1/2 MS medium with or without NaCl, and subsequently subjected for germination in long days at ~22°C. For stratification treatment, seeds on 1/2 MS were kept at 4°C for three days before the germination test. Radicle protrusion is considered as the completion of seed germination. For GA, fluridone and IAA treatments, seeds were sown on 1/2MS medium supplemented with or without the reagents. All germination experiments were performed with three biological replicates, at least 50 seeds were scored for each replicate.

To test the ability of imbibed seeds returning to a dormant state and germinating again, after-ripened seeds imbibed for different times in long days at ~22°C were collected onto a filter paper and dried and stored for one week at room temperature, before subjected to germination again. The germination rates were measured 5 days after second time imbibition with three biological replicates.

### Salt treatment on root growth

For root growth assay, 5-day-old seedlings germinated on 1/2MS medium were transferred onto 1/2MS medium containing NaCl, and seedlings were grown vertically for another 10 days before primary root length measurement. At least 10 seedlings were measured for each line and condition.

### RNA-seq and gene expression analysis

For RNA-seq analysis, after-ripened Col and *ref6*^*c*^ seeds collected at different imbibition time points were subjected for total RNA extraction using Minibest plant RNA extraction kit (Takara). Three independent biological replicates were performed. Sequencing libraries were prepared with BGISEQ-500 RNA-seq library preparation kit according to the manufacture’s instruction. Prepared libraries were sequenced on an MGISEQ-2000 platform and paired-end 150bp reads were generated. Adapter trimming was performed and low-quality reads were filtered with fastp version 0.20.1 (Chen et al., 2018). Reads were mapped to the *Arabidopsis* genome (TAIR10) using Hisat2 version 2.1.10 (Kim et al., 2019). Reads per gene were counted by HTseq version 0.11.2 (Anders et al., 2015). Transcripts per million (TPM) values were generated using R. Differential gene expression analysis was performed using DESeq2 version 1.26.0 (Love et al., 2014). Genes displayed a more than two-fold expression change and had a padj value<0.05 were considered as differentially expressed. Gene ontology analysis was performed with DAVID (https://david.ncifcrf.gov) (Huang et al., 2009).

The expression of *REF6, ELF6, JMJ13* and PRC2 subunits during embryogenesis was extracted from the published datasets (Schneider et al., 2016) with the *Arabidopsis* RNA-seq database (ARS) platform (Zhang et al., 2020). The expression levels were calculated in fragments per kilo base per million mapped reads (FPKM) from three biological replicates.

### RT-qPCR

To analyze the transcripts levels of *GA20ox1, GA20ox2, CYP707A1, CYP707A3, YUC3* and *AUX1*, total RNA was extracted from imbibed seeds with Minibest plant RNA extraction kit (Takara). Reverse transcription was performed using TransScript one-step gDNA removal and cDNA synthesis supermix (TransGen Biotec). Real-time quantitative PCR was conducted on an Applied Biosystems QuantStudio 6 Flex Real-Time PCR System using TransStart top green qPCR supermix (TransGen Biotec). *PP2A* was used as an endogenous control for normalization. Three independent biological replicates were performed for each line and condition. Primers used for amplification are listed in Supplemental Table 1.

### ChIP-seq

ChIP was performed with seeds imbibed for different times or 10-day-old seedlings. Materials were ground with liquid nitrogen into fine powder and fixed with 1% formaldehyde. To profile H3K27me3, nuclei were extracted and mononucleosomes were generated with micrococcal nuclease (MNase) (Sigma, N5386) digestion as previously described (Jiang and Berger, 2017). Immunoprecipitation was conducted with anti-H3K27me3 antibody (Merck, 07-449). For REF6-HA binding ChIP, chromatin was resuspended with ChIP lysis buffer (50mM HEPES (pH7.5), 150mM NaCl, 1mM EDTA, 1% Triton X-100, 0.1% deoxycholate, 0.5% SDS, and 1×protease inhibitor cocktail (Sigma, S8830)) and sheared to generate 200-500bp fragments with Bioruptor (Diagenode) for 30 cycles (30s/on, 30s/off) at high power. Sheared chromatin was diluted four times with ChIP dilution buffer (50mM HEPES (pH7.5), 150mM NaCl, 1mM EDTA, 1% Triton X-100, 0.1% deoxycholate, 0.01% SDS, and 1×protease inhibitor cocktail (Sigma, S8830)) and subjected to immunoprecipitation with anti-HA antibody (CST, 3724). After antibody incubation, Protein A Dynabeads (Life technologies, 10002D) were added to collect the immunocomplexes. After washing, elution, reverse cross-linking and DNA purification, recovered DNA was subjected to library preparation with VAHTS universal DNA library prep kit for illumina (Vazyme, ND607) according to the manufacture’s instruction and sequenced with Illumina NovaSeq 6000 to generate paired-end 150bp reads. All ChIP-seq profiles were performed with two independent biological replicates except for the negative control using Col seedlings in anti-HA ChIP, which was performed with one replicate.

### ChIP-seq data analysis

Adapter trimming was performed and low-quality reads were filtered with fastp version 0.20.1 (Chen et al., 2018). Reads were mapped to the *Arabidopsis* genome (TAIR10) with Botiew2 version 2.4.2 (Langmead and Salzberg, 2012), and filtered for duplicated reads by using Picard version 2.24.0 MarkDuplicates (https://github.com/broadinstitute/picard). The correlation coefficients between two biological replicates were examined using deepTools version 3.1.3 utility multiBamSummary (Supplemental Table 2) (Ramirez et al., 2014), and subsequently two biological replicates were merged with SAMtools version 1.9 (Li et al., 2009). For data visualization, bigwig coverage files were generated using deepTools utility bamCoverage (Ramirez et al., 2014) with a bin size of 10bp and normalized to sequencing depth using reads per kilo base per million mapped reads (RPKM). Visualization was performed with IGV version 2.7.2 (Thorvaldsdottir et al., 2013). Heatmaps and average normalized ChIP-seq profiles were generated using deepTools utilities plotHeatmap and plotProfile. Published ChIP-seq datasets were processed in the same manner (Cui et al., 2016).

ChIP-seq peaks were identified using MACS2 version 2.1.2 with default parameters (Zhang et al., 2008). The parameter ‘--broad’ was used for the calling of H3K27me3 peaks, and narrow peaks were called for REF6-HA ChIP-seq data. The q-value cutoff for peak calling was 0.05. The average scores for H3K27me3 peak regions were calculated with deepTools utility multiBigwigSummary, and differential peaks were called by requiring more than two-fold difference. Genes were assigned to the peaks when the genic regions overlapped with the peak regions for at least one base pair. The identification of DNA motifs underlying REF6 binding peaks were performed as previously described (Cui et al., 2016). Briefly, 800bp genomic regions centered on the REF6 binding peak summits were subjected for motif discovery by MEME-ChIP with default parameters.

### Statistical analysis

Statistical tests performed are indicated in the figure captions. The significance of differences was determined with two-tailed Student’s *t*-test, hypergeometric test, Mann-Whitney U test, and one-way ANOVA with Tukey’s test.

### Accessions numbers

*REF6* (AT3G48430), *ELF6* (AT5G04240), *JMJ13* (AT5G46910), *CLF* (AT2G23380), *SWN* (AT4G02020), *MEA* (AT1G02580), *EMF2* (AT5G51230), *VRN2* (AT4G16845), *FIS2* (AT2G35670), *FIE* (AT3G20740), *DOG1* (AT5G45830), *XTH31* (AT3G44990), *GA20ox1* (AT4G25420), *GA20ox2* (AT5G51810), *CYP707A1* (AT4G19230), *CYP707A3* (AT5G45340), *YUC3* (AT1G04610), *AUX1* (AT2G38120)

## Data availability

The datasets generated during the current study are available in the GEO under accession number GSE167508.

## Supplemental data

**Supplemental Figure 1**. Germination of *ref6* mutants is hypersensitive to NaCl.

**Supplemental Figure 2**. H3K27me3 changes at early phase germination.

**Supplemental Figure 3**. H3K27me3 changes at late phase germination.

**Supplemental Figure 4**. GO analysis of H3K27me3 increased and decreased genes at late phase germination.

**Supplemental Figure 5**. Differential H3K27me3 analysis in Col and *ref6* mutants.

**Supplemental Figure 6**. Overlap analysis between hypermethylated genes and expression down-regulated genes in *ref6*^*c*^.

**Supplemental Figure 7**. Genomic binding of REF6-HA.

**Supplemental Figure 8**. REF6 demethylates H3K27 at GA, ABA and auxin-related genes.

**Supplemental Figure 9**. Expression of cell cycle genes during germination.

**Supplemental Table 1**. RT-qPCR primers used in this study.

**Supplemental Table 2**. The correlation coefficients between ChIP-seq replicates.

**Supplemental Dataset 1**. Lists of mis-expressed genes in *ref6*^*c*^ compared with Col during germination.

**Supplemental Dataset 2**. Lists of differential H3K27me3 marked genes in 6h, 24h, 48h, and 72h imbibed Col seeds compared with 0h imbibed seeds determined by ChIP-seq.

**Supplemental Dataset 3**. Lists of genes with increased and decreased H3K27me3 in *ref6*^*c*^ compared with Col during germination.

**Supplemental Dataset 4**. Lists of REF6-HA binding peaks in 0h, 6h, 24h, and 48h imbibed seeds and 10-day-old seedlings.

## Authors’ contributions

D.J. conceived the research. J.P., Z.Z., and T.Z. performed experiments. J.P., H.Z., and D.J. performed bioinformatics analysis. J.P., H.Z., and D.J. interpreted the data. D.J. wrote the paper.

## Acknowledgments

We thank Dr. Xiaofeng Cao for kindly providing seeds, Drs. Xiaofeng Cao and Mike Borg for critical reading of the manuscript and valuable suggestions.

## References

Anders S, Pyl PT, Huber W (2015) HTSeq-a Python framework to work with high-throughput sequencing data. Bioinformatics 31: 166–169

Antunez-Sanchez J, Naish M, Ramirez-Prado JS, Ohno S, Huang Y, Dawson A, Opassathian K, Manza-Mianza D, Ariel F, Raynaud C, Wibowo A, Daron J, Ueda M, Latrasse D, Slotkin RK, Weigel D, Benhamed M, Gutierrez-Marcos J (2020) A new role for histone demethylases in the maintenance of plant genome integrity. Elife 9

Barroco RM, Van Poucke K, Bergervoet JHW, De Veylder L, Groot SPC, Inze D, Engler G (2005) The role of the cell cycle machinery in resumption of postembryonic development. Plant Physiology 137: 127–140

Borg M, Jacob Y, Susaki D, LeBlanc C, Buendia D, Axelsson E, Kawashima T, Voigt P, Boavida L, Becker J, Higashiyama T, Martienssen R, Berger F (2020) Targeted reprogramming of H3K27me3 resets epigenetic memory in plant paternal chromatin. Nature Cell Biology 22: 621-+

Bouyer D, Roudier F, Heese M, Andersen ED, Gey D, Nowack MK, Goodrich J, Renou JP, Grini PE, Colot V, Schnittger A (2011) Polycomb Repressive Complex 2 Controls the Embryo-to-Seedling Phase Transition. Plos Genetics 7

Brady SM, Sarkar SF, Bonetta D, McCourt P (2003) The ABSCISIC ACID INSENSITIVE 3 (ABI3) gene is modulated by farnesylation and is involved in auxin signaling and lateral root development in Arabidopsis. Plant Journal 34: 67–75

Chanvivattana Y, Bishopp A, Schubert D, Stock C, Moon YH, Sung ZR, Goodrich J (2004) Interaction of polycomb-group proteins controlling flowering in Arabidopsis. Development 131: 5263–5276

Chen H, Tong J, Fu W, Liang Z, Ruan J, Yu Y, Song X, Yuan L, Xiao L, Liu J, Cui Y, Huang S, Li C (2020) The H3K27me3 Demethylase RELATIVE OF EARLY FLOWERING6 Suppresses Seed Dormancy by Inducing Abscisic Acid Catabolism. Plant Physiol 184: 1969–1978

Chen SF, Zhou YQ, Chen YR, Gu J (2018) fastp: an ultra-fast all-in-one FASTQ preprocessor. Bioinformatics 34: 884–890

Coleman RT, Struhl G (2017) Causal role for inheritance of H3K27me3 in maintaining the OFF state of a Drosophila HOX gene. Science 356

Comai L, Harada JJ (1990) Transcriptional Activities in Dry Seed Nuclei Indicate the Timing of the Transition from Embryogeny to Germination. Proceedings of the National Academy of Sciences of the United States of America 87: 2671–2674

Corbineau F, Xia Q, Bailly C, El-Maarouf-Bouteau H (2014) Ethylene, a key factor in the regulation of seed dormancy. Frontiers in Plant Science 5

Cosgrove DJ (2015) Plant expansins: diversity and interactions with plant cell walls. Current Opinion in Plant Biology 25: 162–172

Crevillen P, Yang H, Cui X, Greeff C, Trick M, Qiu Q, Cao X, Dean C (2014) Epigenetic reprogramming that prevents transgenerational inheritance of the vernalized state. Nature 515: 587–590

Cui X, Lu F, Qiu Q, Zhou B, Gu L, Zhang S, Kang Y, Cui X, Ma X, Yao Q, Ma J, Zhang X, Cao X (2016) REF6 recognizes a specific DNA sequence to demethylate H3K27me3 and regulate organ boundary formation in Arabidopsis. Nat Genet 48: 694–699

Finkelstein R, Reeves W, Ariizumi T, Steber C (2008) Molecular aspects of seed dormancy. Annual Review of Plant Biology 59: 387–415

Finkelstein RR, Gampala SS, Rock CD (2002) Abscisic acid signaling in seeds and seedlings. Plant Cell 14 Suppl: S15-45

Gan ES, Xu YF, Wong JY, Goh JG, Sun B, Wee WY, Huang JB, Ito T (2014) Jumonji demethylases moderate precocious flowering at elevated temperature via regulation of FLC in Arabidopsis. Nature Communications 5

Grossniklaus U, Vielle-Calzada JP, Hoeppner MA, Gagliano WB (1998) Maternal control of embryogenesis by medea, a Polycomb group gene in Arabidopsis. Science 280: 446–450

Gu XF, Xu TD, He YH (2014) A Histone H3 Lysine-27 Methyltransferase Complex Represses Lateral Root Formation in Arabidopsis thaliana. Molecular Plant 7: 977–988

He JN, Duan Y, Hua DP, Fan GJ, Wang L, Liu Y, Chen ZZ, Han LH, Qu LJ, Gong ZZ (2012) DEXH Box RNA Helicase-Mediated Mitochondrial Reactive Oxygen Species Production in Arabidopsis Mediates Crosstalk between Abscisic Acid and Auxin Signaling. Plant Cell 24: 1815–1833

Hepworth J, Dean C (2015) Flowering Locus C’s Lessons: Conserved Chromatin Switches Underpinning Developmental Timing and Adaptation. Plant Physiology 168: 1237–1245

Holdsworth MJ, Bentsink L, Soppe WJJ (2008) Molecular networks regulating Arabidopsis seed maturation, after-ripening, dormancy and germination. New Phytologist 179: 33–54

Hosogane M, Funayama R, Shirota M, Nakayama K (2016) Lack of Transcription Triggers H3K27me3 Accumulation in the Gene Body. Cell Reports 16: 696–706

Hou XL, Zhou JN, Liu C, Liu L, Shen LS, Yu H (2014) Nuclear factor Y-mediated H3K27me3 demethylation of the SOC1 locus orchestrates flowering responses of Arabidopsis. Nature Communications 5

Hsueh YL, Lou CH (1947) Effects of 2,4-D on Seed Germination and Respiration. Science 105: 283–285

Huang DW, Sherman BT, Lempicki RA (2009) Systematic and integrative analysis of large gene lists using DAVID bioinformatics resources. Nature Protocols 4: 44–57

Hyun Y, Richter R, Vincent C, Martinez-Gallegos R, Porri A, Coupland G (2016) Multi-layered Regulation of SPL15 and Cooperation with SOC1 Integrate Endogenous Flowering Pathways at the Arabidopsis Shoot Meristem. Developmental Cell 37: 254–266

Ikeuchi M, Iwase A, Rymen B, Harashima H, Shibata M, Ohnuma M, Breuer C, Morao AK, de Lucas M, De Veylder L, Goodrich J, Brady SM, Roudier F, Sugimoto K (2015) PRC2 represses dedifferentiation of mature somatic cells in Arabidopsis. Nat Plants 1: 15089

Jiang D, Berger F (2017) DNA replication-coupled histone modification maintains Polycomb gene silencing in plants. Science 357: 1146–1149

Jiang D, Wang Y, Wang Y, He Y (2008) Repression of FLOWERING LOCUS C and FLOWERING LOCUS T by the Arabidopsis Polycomb repressive complex 2 components. PLoS One 3: e3404

Jurgens G (1985) A Group of Genes-Controlling the Spatial Expression of the Bithorax Complex in Drosophila. Nature 316: 153–155

Katz A, Oliva M, Mosquna A, Hakim O, Ohad N (2004) FIE and CURLY LEAF polycomb proteins interact in the regulation of homeobox gene expression during sporophyte development. Plant Journal 37: 707–719

Kermode AR (2005) Role of abscisic acid in seed dormancy. Journal of Plant Growth Regulation 24: 319–344

Kim D, Paggi JM, Park C, Bennett C, Salzberg SL (2019) Graph-based genome alignment and genotyping with HISAT2 and HISAT-genotype. Nature Biotechnology 37: 907-+

Kim DH, Doyle MR, Sung S, Amasino RM (2009) Vernalization: winter and the timing of flowering in plants. Annu Rev Cell Dev Biol 25: 277–299

Lafos M, Kroll P, Hohenstatt ML, Thorpe FL, Clarenz O, Schubert D (2011) Dynamic regulation of H3K27 trimethylation during Arabidopsis differentiation. PLoS Genet 7: e1002040

Langmead B, Salzberg SL (2012) Fast gapped-read alignment with Bowtie 2. Nature Methods 9: 357–U354

Lee LR, Wengier DL, Bergmann DC (2019) Cell-type-specific transcriptome and histone modification dynamics during cellular reprogramming in the Arabidopsis stomatal lineage (vol 116, pg 21914, 2019). Proceedings of the National Academy of Sciences of the United States of America 116: 24376–24376

Li C, Gu L, Gao L, Chen C, Wei CQ, Qiu Q, Chien CW, Wang S, Jiang L, Ai LF, Chen CY, Yang S, Nguyen V, Qi Y, Snyder MP, Burlingame AL, Kohalmi SE, Huang S, Cao X, Wang ZY, Wu K, Chen X, Cui Y (2016) Concerted genomic targeting of H3K27 demethylase REF6 and chromatin-remodeling ATPase BRM in Arabidopsis. Nat Genet 48: 687–693

Li H, Handsaker B, Wysoker A, Fennell T, Ruan J, Homer N, Marth G, Abecasis G, Durbin R, Proc GPD (2009) The Sequence Alignment/Map format and SAMtools. Bioinformatics 25: 2078–2079

Li Z, Ou Y, Zhang Z, Li J, He Y (2018) Brassinosteroid Signaling Recruits Histone 3 Lysine-27 Demethylation Activity to FLOWERING LOCUS C Chromatin to Inhibit the Floral Transition in Arabidopsis. Mol Plant 11: 1135–1146

Liu C, Lu F, Cui X, Cao X (2010) Histone methylation in higher plants. Annu Rev Plant Biol 61: 395–420

Liu PP, Montgomery TA, Fahlgren N, Kasschau KD, Nonogaki H, Carrington JC (2007) Repression of AUXIN RESPONSE FACTOR10 by microRNA160 is critical for seed germination and post-germination stages. Plant Journal 52: 133–146

Liu XG, Kim YJ, Muller R, Yumul RE, Liu CY, Pan YY, Cao XF, Goodrich J, Chena XM (2011) AGAMOUS Terminates Floral Stem Cell Maintenance in Arabidopsis by Directly Repressing WUSCHEL through Recruitment of Polycomb Group Proteins. Plant Cell 23: 3654–3670

Liu Y, Koornneef M, Soppe WJ (2007) The absence of histone H2B monoubiquitination in the Arabidopsis hub1 (rdo4) mutant reveals a role for chromatin remodeling in seed dormancy. Plant Cell 19: 433–444

Love MI, Huber W, Anders S (2014) Moderated estimation of fold change and dispersion for RNA-seq data with DESeq2. Genome Biology 15

Lu F, Cui X, Zhang S, Jenuwein T, Cao X (2011) Arabidopsis REF6 is a histone H3 lysine 27 demethylase. Nat Genet 43: 715–719

Luo X, He YH (2020) Experiencing winter for spring flowering: A molecular epigenetic perspective on vernalization. Journal of Integrative Plant Biology 62: 104–117

Marowa P, Ding AM, Kong YZ (2016) Expansins: roles in plant growth and potential applications in crop improvement. Plant Cell Reports 35: 949–965

Molitor AM, Bu Z, Yu Y, Shen WH (2014) Arabidopsis AL PHD-PRC1 complexes promote seed germination through H3K4me3-to-H3K27me3 chromatin state switch in repression of seed developmental genes. PLoS Genet 10: e1004091

Mozgova I, Hennig L (2015) The Polycomb Group Protein Regulatory Network. Annual Review of Plant Biology, Vol 66 66: 269–296

Mozgova I, Munoz-Viana R, Hennig L (2017) PRC2 Represses Hormone-Induced Somatic Embryogenesis in Vegetative Tissue of Arabidopsis thaliana. PLoS Genet 13: e1006562

Nee G, Xiang Y, Soppe WJJ (2017) The release of dormancy, a wake-up call for seeds to germinate. Current Opinion in Plant Biology 35: 8–14

Noh B, Lee SH, Kim HJ, Yi G, Shin EA, Lee M, Jung KJ, Doyle MR, Amasino RM, Noh YS (2004) Divergent roles of a pair of homologous jumonji/zinc-finger-class transcription factor proteins in the regulation of Arabidopsis flowering time. Plant Cell 16: 2601–2613

Nonogaki H (2014) Seed dormancy and germination-emerging mechanisms and new hypotheses. Front Plant Sci 5: 233

Ogawa M, Hanada A, Yamauchi Y, Kuwalhara A, Kamiya Y, Yamaguchi S (2003) Gibberellin biosynthesis and response during Arabidopsis seed germination. Plant Cell 15: 1591–1604

Okamoto M, Kuwahara A, Seo M, Kushiro T, Asami T, Hirai N, Kamiya Y, Koshiba T, Nambara E (2006) CYP707A1 and CYP707A2, which encode abscisic acid 8 ‘-hydroxylases, are indispensable for proper control of seed dormancy and germination in Arabidopsis. Plant Physiology 141: 97–107

Qiu Q, Mei H, Deng X, He K, Wu B, Yao Q, Zhang J, Lu F, Ma J, Cao X (2019) DNA methylation repels targeting of Arabidopsis REF6. Nat Commun 10: 2063

Ramirez F, Dundar F, Diehl S, Gruning BA, Manke T (2014) deepTools: a flexible platform for exploring deep-sequencing data. Nucleic Acids Research 42: W187–W191

Riising EM, Comet I, Leblanc B, Wu XD, Johansen JV, Helin K (2014) Gene Silencing Triggers Polycomb Repressive Complex 2 Recruitment to CpG Islands Genome Wide. Molecular Cell 55: 347–360

Schneider A, Aghamirzaie D, Elmarakeby H, Poudel AN, Koo AJ, Heath LS, Grene R, Collakova E (2016) Potential targets of VIVIPAROUS1/ABI3-LIKE1 (VAL1) repression in developing Arabidopsis thaliana embryos. Plant Journal 85: 305–319

Smaczniak C, Immink RGH, Muino JM, Blanvillain R, Busscher M, Busscher-Lange J, Dinh QD, Liu SJ, Westphal AH, Boeren S, Parcy F, Xu L, Carles CC, Angenent GC, Kaufmann K (2012) Characterization of MADS-domain transcription factor complexes in Arabidopsis flower development. Proceedings of the National Academy of Sciences of the United States of America 109: 1560–1565

Steber CM, McCourt P (2001) A role for brassinosteroids in germination in Arabidopsis. Plant Physiology 125: 763–769

Sun B, Looi LS, Guo SY, He ZM, Gan ES, Huang JB, Xu YF, Wee WY, Ito T (2014) Timing Mechanism Dependent on Cell Division Is Invoked by Polycomb Eviction in Plant Stem Cells. Science 343: 505-+

Thorvaldsdottir H, Robinson JT, Mesirov JP (2013) Integrative Genomics Viewer (IGV): high-performance genomics data visualization and exploration. Briefings in Bioinformatics 14: 178–192

van Zanten M, Koini MA, Geyer R, Liu Y, Brambilla V, Bartels D, Koornneef M, Fransz P, Soppe WJ (2011) Seed maturation in Arabidopsis thaliana is characterized by nuclear size reduction and increased chromatin condensation. Proc Natl Acad Sci U S A 108: 20219–20224

van Zanten M, Zoll C, Wang Z, Philipp C, Carles A, Li Y, Kornet NG, Liu Y, Soppe WJ (2014) HISTONE DEACETYLASE 9 represses seedling traits in Arabidopsis thaliana dry seeds. Plant J 80: 475–488

Wang XL, Gao J, Gao S, Li ZP, Kuai BK, Ren GD (2019) REF6 promotes lateral root formation through de-repression of PIN1/3/7 genes. Journal of Integrative Plant Biology 61: 383–387

Wang XL, Gao J, Gao S, Song Y, Yang Z, Kuai BK (2019) The H3K27me3 demethylase REF6 promotes leaf senescence through directly activating major senescence regulatory and functional genes in Arabidopsis. Plos Genetics 15

Wang Z, Chen FY, Li XY, Cao H, Ding M, Zhang C, Zuo JH, Xu CN, Xu JM, Deng X, Xiang Y, Soppe WJJ, Liu YX (2016) Arabidopsis seed germination speed is controlled by SNL histone deacetylase-binding factor-mediated regulation of AUX1. Nature Communications 7

Wiles ET, Selker EU (2017) H3K27 methylation: a promiscuous repressive chromatin mark. Current Opinion in Genetics & Development 43: 31–37

Yan W, Chen D, Smaczniak C, Engelhorn J, Liu H, Yang W, Graf A, Carles CC, Zhou DX, Kaufmann K (2018) Dynamic and spatial restriction of Polycomb activity by plant histone demethylases. Nat Plants 4: 681–689

Yu XF, Li L, Li L, Guo M, Chory J, Yin YH (2008) Modulation of brassinosteroid-regulated gene expression by jumonji domain-containing proteins ELF6 and REF6 in Arabidopsis. Proceedings of the National Academy of Sciences of the United States of America 105: 7618–7623

Zander M, Willige BC, He YP, Nguyen TA, Langford AE, Nehring R, Howell E, McGrath R, Bartlett A, Castanon R, Nery JR, Chen HM, Zhang ZZ, Jupe F, Stepanova A, Schmitz RJ, Lewsey MG, Chory J, Ecker JR (2019) Epigenetic silencing of a multifunctional plant stress regulator. Elife 8

Zhang H, Zhang F, Yu YM, Feng L, Jia JB, Liu B, Li BS, Guo HW, Zhai JX (2020) A Comprehensive Online Database for Exploring similar to 20,000 Public Arabidopsis RNA-Seq Libraries. Molecular Plant 13: 1231–1233

Zhang X, Clarenz O, Cokus S, Bernatavichute YV, Pellegrini M, Goodrich J, Jacobsen SE (2007) Whole-genome analysis of histone H3 lysine 27 trimethylation in Arabidopsis. PLoS Biol 5: e129

Zhang Y, Liu T, Meyer CA, Eeckhoute J, Johnson DS, Bernstein BE, Nusbaum C, Myers RM, Brown M, Li W, Liu XS (2008) Model-based analysis of ChIP-Seq (MACS). Genome Biol 9: R137

Zhao M, Yang S, Liu X, Wu K (2015) Arabidopsis histone demethylases LDL1 and LDL2 control primary seed dormancy by regulating DELAY OF GERMINATION 1 and ABA signaling-related genes. Front Plant Sci 6: 159

Zheng J, Chen F, Wang Z, Cao H, Li X, Deng X, Soppe WJ, Li Y, Liu Y (2012) A novel role for histone methyltransferase KYP/SUVH4 in the control of Arabidopsis primary seed dormancy. New Phytol 193: 605–616

Zheng S, Hu H, Ren H, Yang Z, Qiu Q, Qi W, Liu X, Chen X, Cui X, Li S, Zhou B, Sun D, Cao X, Du J (2019) The Arabidopsis H3K27me3 demethylase JUMONJI 13 is a temperature and photoperiod dependent flowering repressor. Nat Commun 10: 1303

