## Supplemental Figures for "The REF6-dependent H3K27 demethylation establishes transcriptional competence to promote germination in *Arabidopsis*"

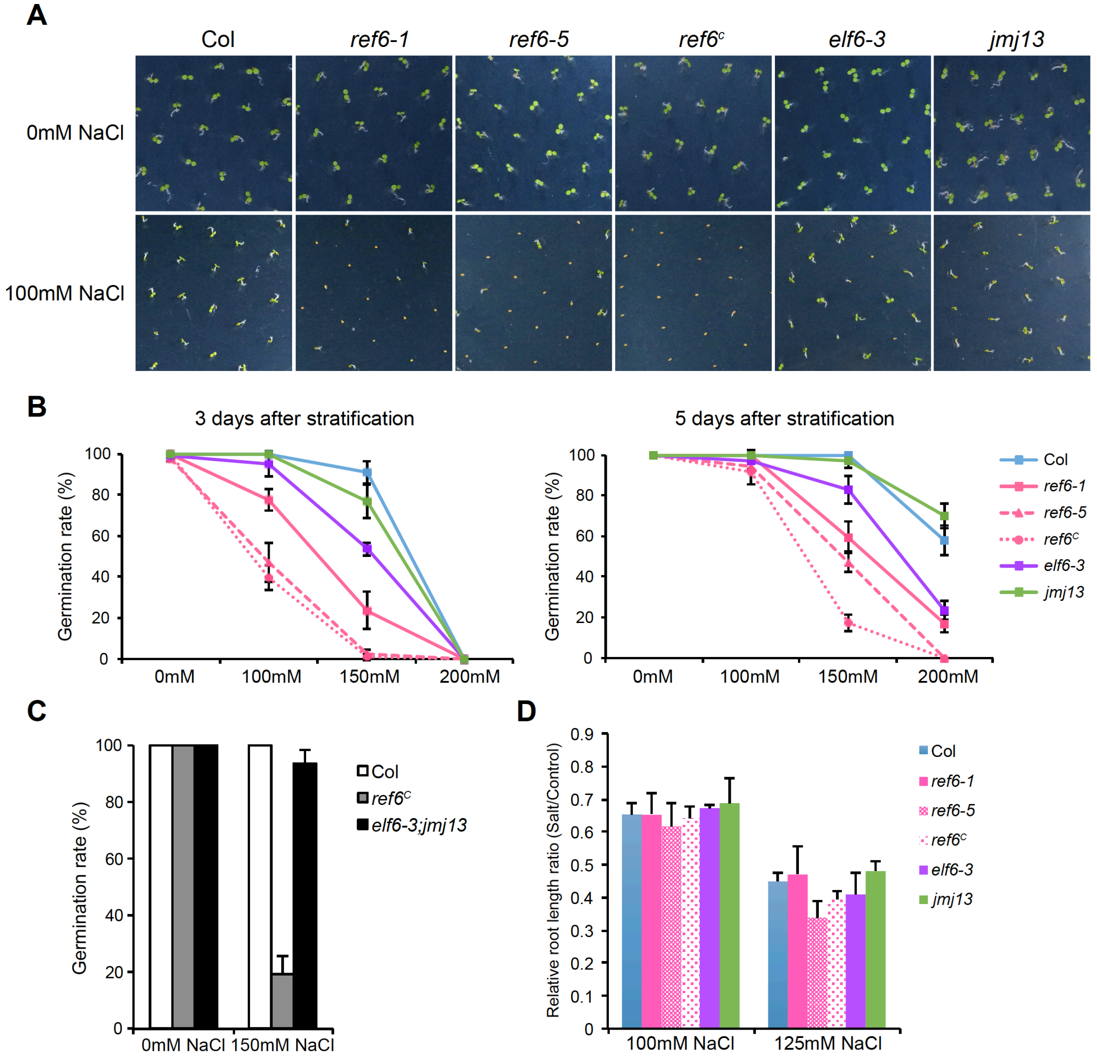


**Supplemental Figure 1. Germination of *ref6* mutants is hypersensitive to NaCl**

**A.** The germination phenotypes of after-ripened and stratified seeds imbibed for 3 days on 1/2MS medium with or without NaCl.

**B.** The germination rates of after-ripened and stratified seeds imbibed on 1/2MS medium with different concentrations of NaCl. Germination rates were measured 3 days and 5 days after stratification treatment. Values are means ± SD of three biological repeats. At least 50 seeds were analyzed for each replicate.

**C.** The germination rates of after-ripened and stratified seeds imbibed on 1/2MS medium with or without NaCl. Germination rates were measured 5 days after stratification treatment. Values are means ± SD of three biological repeats. At least 50 seeds were analyzed for each replicate.

**D.** Relative primary root length ratios of seedlings grown on 1/2MS medium with NaCl compared with each genotype grown on 1/2MS without NaCl, at least 10 seedlings for each line and condition were measured.

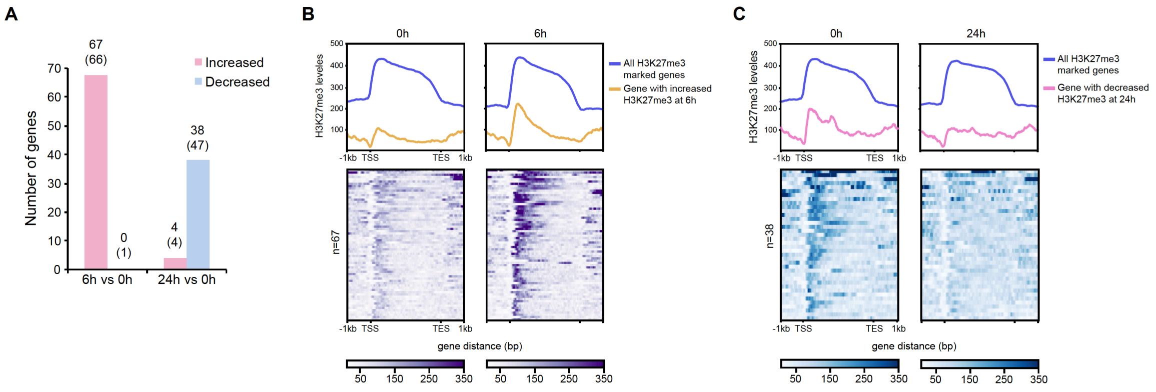


**Supplemental Figure 2. H3K27me3 changes at early phase germination**

**A.** Numbers of H3K27me3 increased and decreased genes in 6h and 24h imbibed Col seeds compared with 0h imbibed seeds determined by ChIP-seq. Numbers in brackets indicate the amount of differential H3K27me3 peaks identified by ChIP-seq.

**B.** Average profiles and heat maps of normalized H3K27me3 ChIP-seq signals over H3K27me3 increased genes in 6h imbibed Col seeds (n=67). The profiles were generated after merging two biological replicates at each time point.

**C.** Average profiles and heat maps of normalized H3K27me3 ChIP-seq signals over H3K27me3 decreased genes in 24h imbibed Col seeds (n=38). The profiles were generated after merging two biological replicates at each time point.

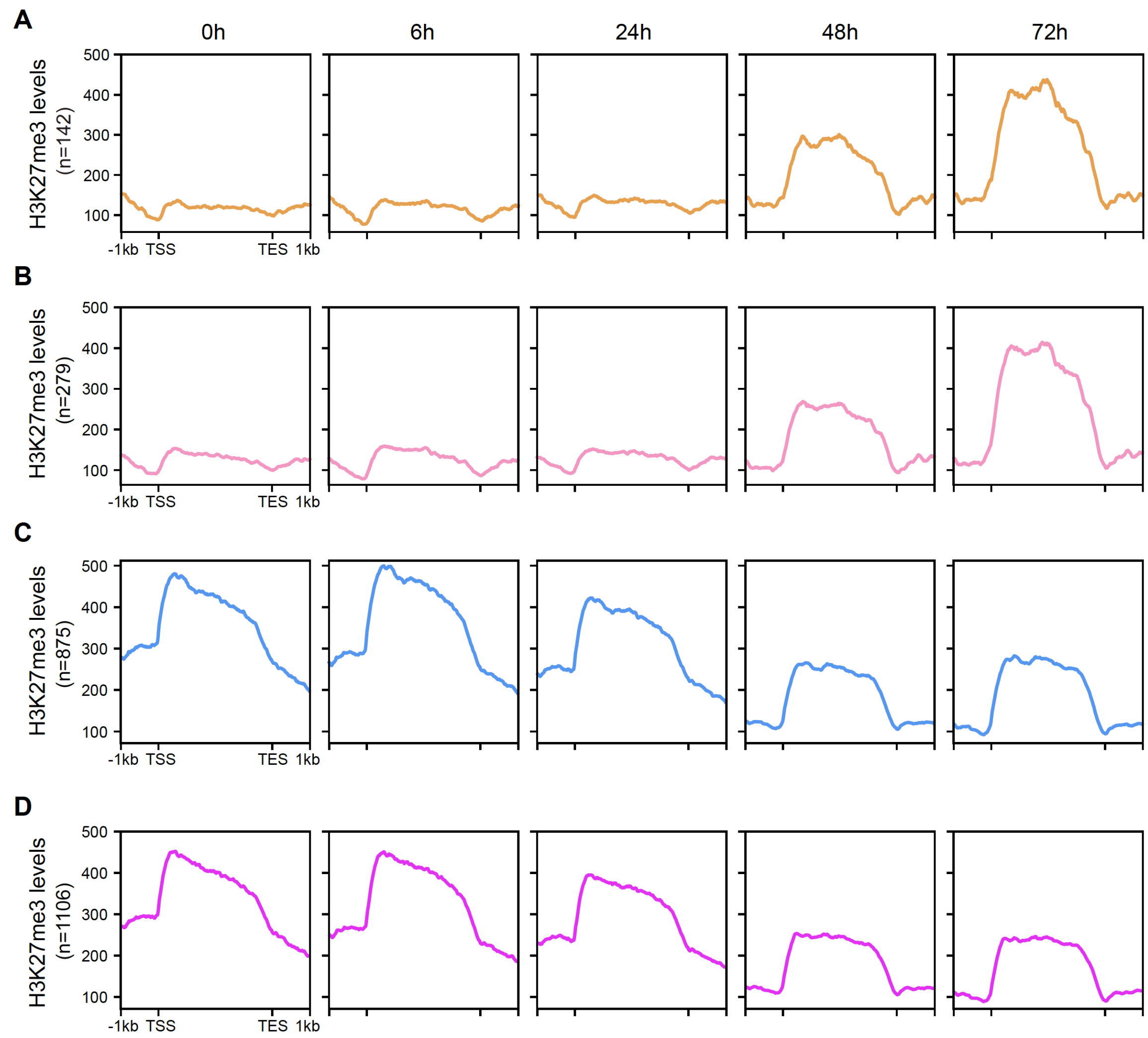


**Supplemental Figure 3. H3K27me3 changes at late phase germination**

**A, B.** Average profiles of normalized H3K27me3 ChIP-seq signals over H3K27me3 increased genes identified in 48h imbibed Col seeds (n=142) (A) and 72h imbibed Col seeds (n=279) (B). The profiles were generated after merging two biological replicates at each time point.

**C, D.** Average profiles of normalized H3K27me3 ChIP-seq signals over H3K27me3 decreased genes identified in 48h imbibed Col seeds (n=875) (C) and 72h imbibed Col seeds (n=1106) (D). The profiles were generated after merging two biological replicates at each time point.

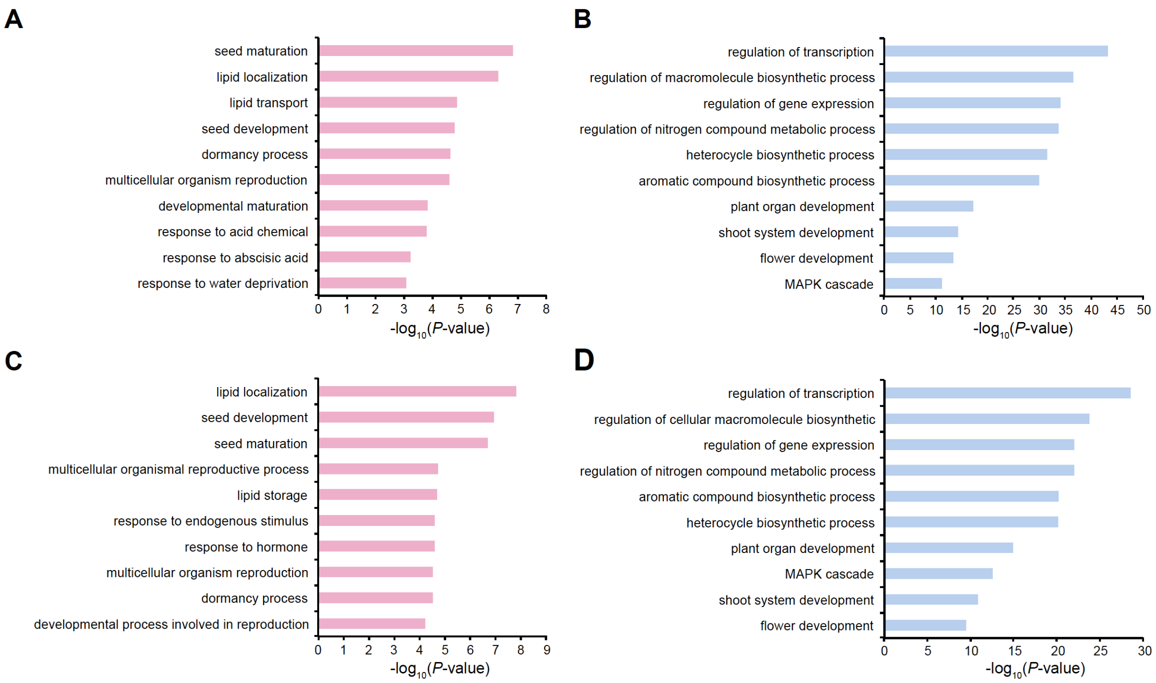


**Supplemental Figure 4. GO analysis of H3K27me3 increased and decreased genes at late phase germination**

**A, B.** GO analysis of H3K27me3 increased (A) and decreased (B) genes in 48h imbibed Col seeds. Top 10 representative terms are listed and ranked by *P* value.

**C, D.** GO analysis of H3K27me3 increased (C) and decreased (D) genes in 72h imbibed Col seeds. Top 10 representative terms are listed and ranked by *P* value.

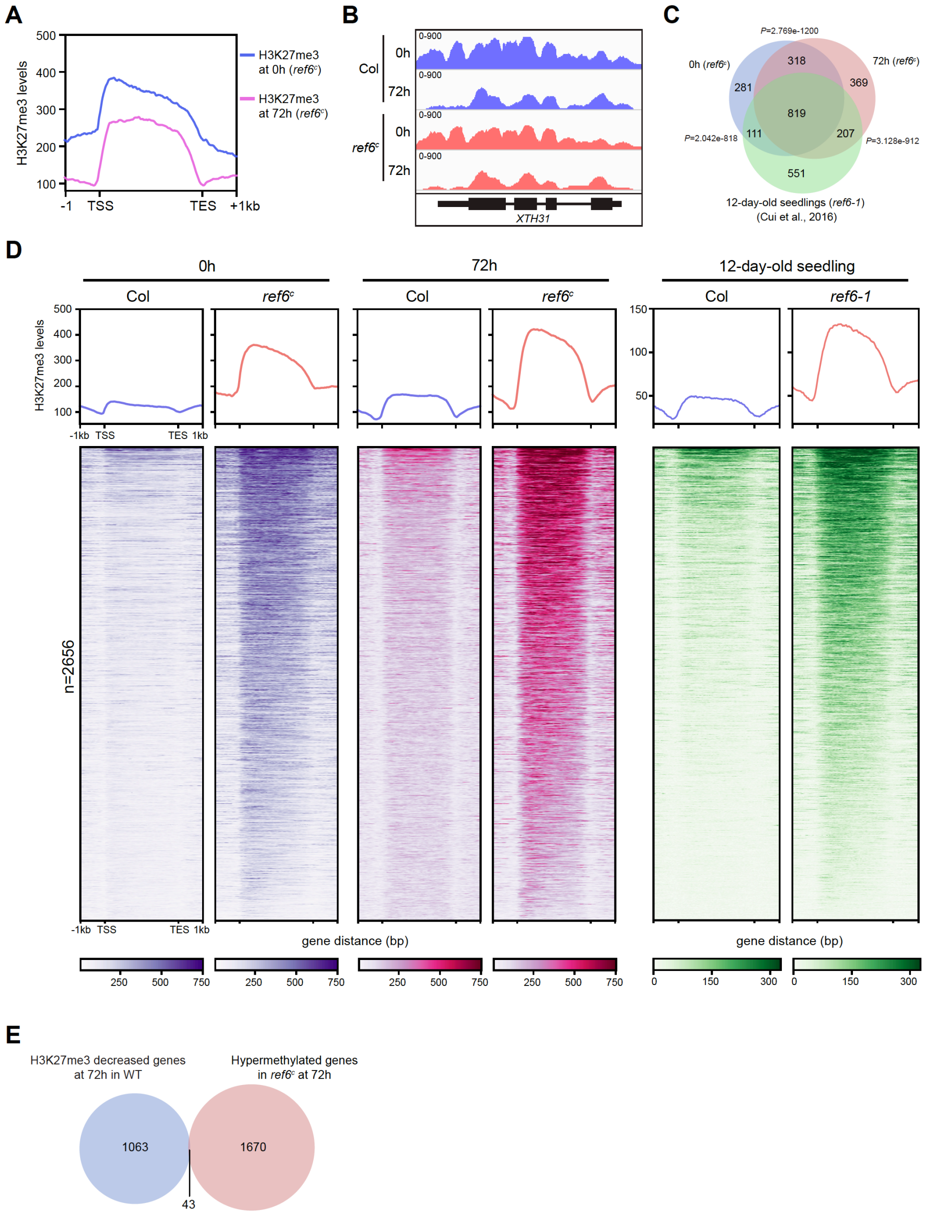


**Supplemental Figure 5. Differential H3K27me3 analysis in Col and *ref6* mutants**

**A.** Average profiles of normalized H3K27me3 ChIP-seq signals in 0h and 72h imbibed *ref6*$^{c}$ seeds over H3K27me3 decreased genes identified in 72h imbibed Col seeds (n=1106). The profiles were generated after merging two biological replicates at each time point.

**B.** Genome browser view of H3K27me3 enrichment at the *XTH31* locus in Col and *ref6*$^{c}$ seeds imbibed for 0h and 72h. The profiles were generated after merging two biological replicates at each time point.

**C.** Venn diagram of H3K27me3 hypermethylated genes in 0h and 72h imbibed *ref6*$^{c}$ seeds and 12-day-old *ref6-1* seedlings (Cui et al., 2016). *P* values are based on the hypergeometric test.

**D.** Average profiles and heat maps of normalized H3K27me3 ChIP-seq signals over a combined set of hypermethylated genes identified in 0h and 72h imbibed *ref6*$^{c}$ seeds and 12-day-old *ref6-1* seedlings. The profiles were generated after merging two biological replicates at each time point.

**E.** Venn diagram of H3K27me3 decreased genes in 72h imbibed Col seeds compared with 0h imbibed seeds and hypermethylated genes in *ref6*$^{c}$ compared with Col. *P* values are based on the hypergeometric test.

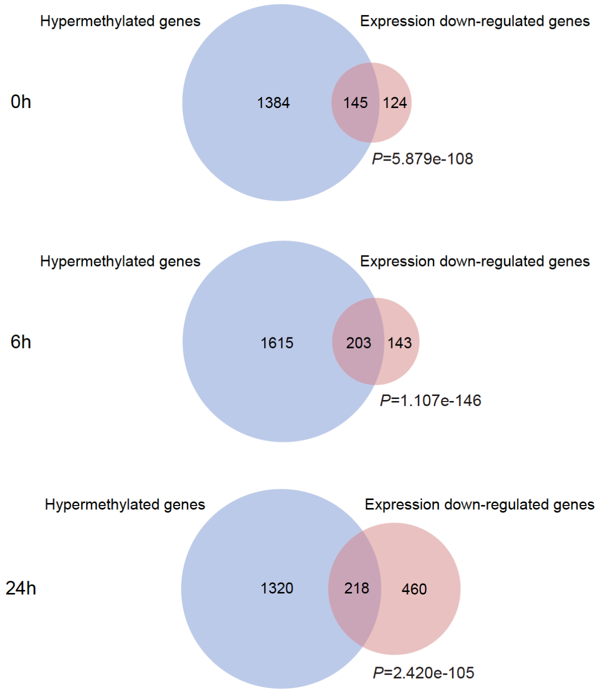


**Supplemental Figure 6. Overlap analysis between hypermethylated genes and expression down-regulated genes in *ref6*$^c$.** Venn diagrams of hypermethylated genes and expression down-regulated genes in *ref6*$^c$ compared with Col at each imbibition time point. *P* values are based on the hypergeometric test.

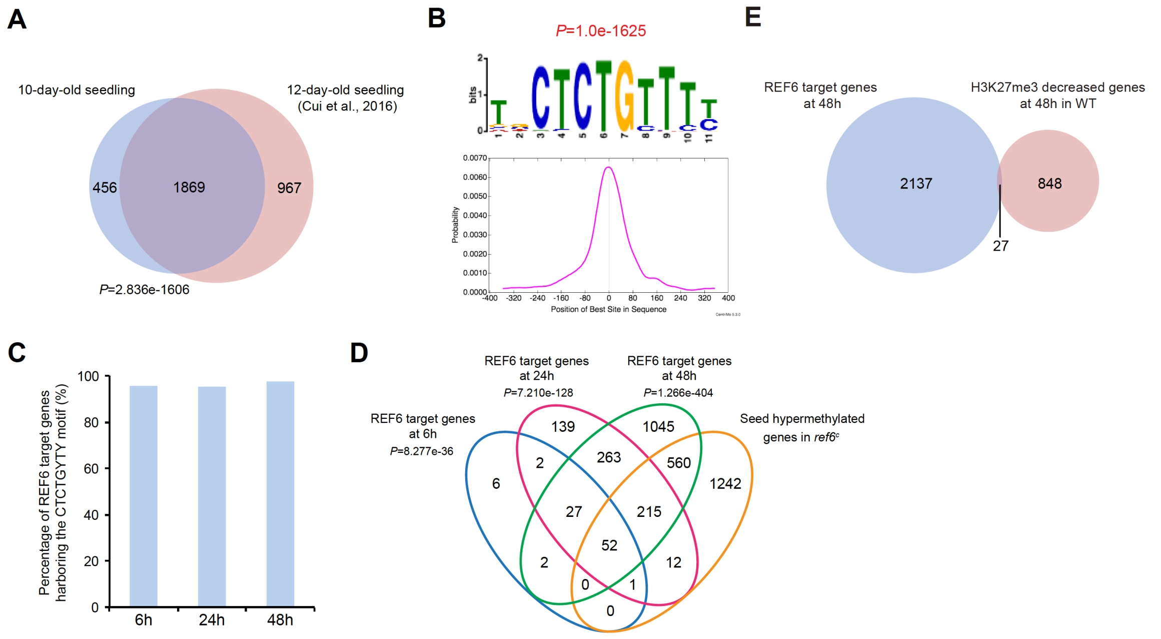


**Supplemental Figure 7. Genomic binding of REF6-HA**

**A.** Venn diagram of REF6 target genes in 10-day-old seedlings identified in this study and those in 12-day-old seedlings identified in a previous study (Cui et al., 2016). *P* values are based on the hypergeometric test.

**B.** The top DNA motif enriched and its distribution in REF6 binding sites in 10-day-old seedlings.

**C.** The percentage of identified REF6 target genes in seeds containing the CTCTGYTY motif.

**D.** Venn diagram of combined seed hypermethylated genes in 0h, 6h and 24h imbibed *ref6*$^{c}$ seeds with REF6 target genes at 6h, 24h and 48h.

**E.** Venn diagram of REF6 target genes in 48h imbibed seeds and H3K27me3 decreased genes in 48h imbibed Col seeds compared with 0h imbibed Col seeds.

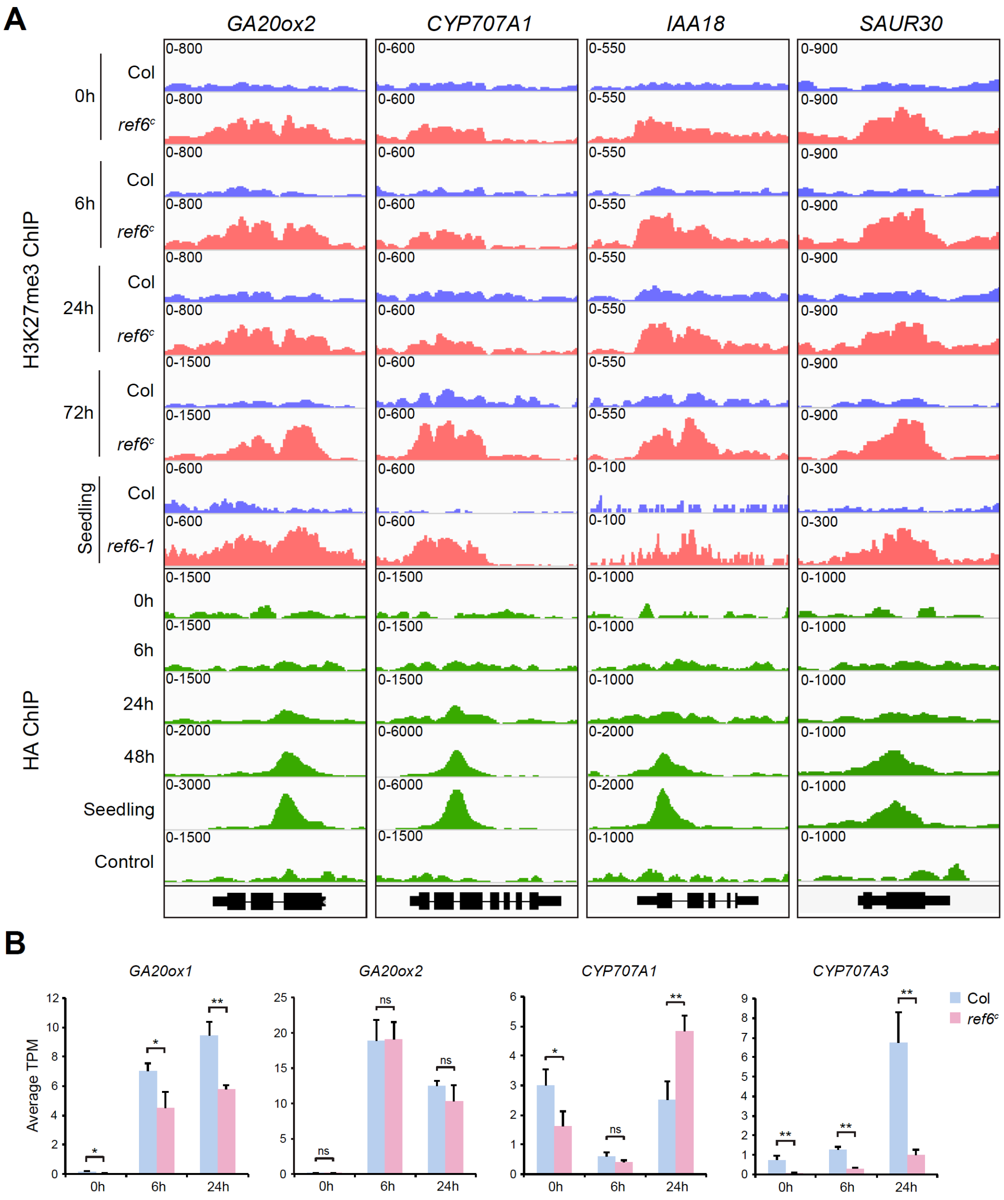


**Supplemental Figure 8. REF6 demethylates H3K27 at GA, ABA and auxin-related genes**

**A.** Genome browser views of H3K27me3 and REF6-HA enrichment at *GA20ox2*, *CYP7070A1*, *IAA18* and *SAUR30* loci in imbibed seeds and seedlings. Col seedlings were used as a negative control for the HA ChIP. The profiles were generated after merging two biological replicates at each time point.

**B.** Expression of *GA20ox1*, *GA20ox2*, *CYP707A1* and *CYP707A3* in Col and *ref6*$^{c}$ during germination determined by RNA-seq. Values are means ± SD of three biological repeats. Statistical significance was determined by two-tailed Student's *t*-test (ns, not significant; *, *P*<0.05; **, *P*<0.01).

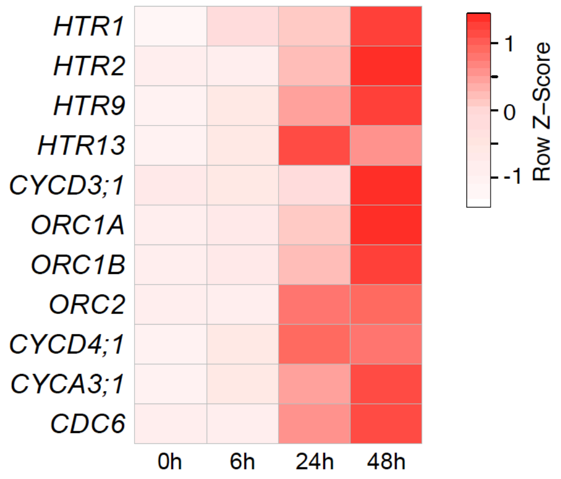


**Supplemental Figure 9. Expression of cell cycle genes during germination**

Heat map showing the relative expression changes (z-normalized) of cell cycle genes during germination in Col. The expression values represent average of three independent RNA-seq replicates.

**References**

**Cui X, Lu F, Qiu Q, Zhou B, Gu L, Zhang S, Kang Y, Cui X, Ma X, Yao Q, Ma J, Zhang X, Cao X** (2016) REF6 recognizes a specific DNA sequence to demethylate H3K27me3 and regulate organ boundary formation in Arabidopsis. Nat Genet **48:** 694-699
