## Supplemental Tables for "The REF6-dependent H3K27 demethylation establishes transcriptional competence to promote germination in *Arabidopsis*"

**Supplemental Table 1. RT-qPCR primers used in this study.**

| Gene | Primer (5' to 3') |
|---|---|
| *GA20ox1* | CGATACTTTCATGGCTCTATCG |
| | ATGCTGTCCAAAAGCTCTCTCG |
| *GA20ox2* | GAGCAGTTTGGGAAGGTGTATC |
| | CGTTCTCTTCGAAAAATCCTCG |
| *CYP707A1* | ATCAAGATTCGAGGTGGCTCC |
| | CTTGGTGGTGAGATGATGAATC |
| *CYP707A3* | ACTAAGTACAGATGGTCAATCG |
| | ATCTATGGTTTTCGTTCCAAGG |
| *YUC3* | ATGGCTTAAGGACAACGACTTC |
| | ACACTCATAGCATCAAGCGACG |
| *AUX1* | TTGTACACCTTTGGAGGTCACG |
| | GCTGACGGAATCGTTAGCGTG |

**Supplemental Table 2. The correlation coefficients between ChIP-seq replicates.**

| Sample | R value |
| --- | --- |
| Col-Input-0h-1_vs_Col-Input-0h-2 | 0.991 |
| Col-Input-6h-1_vs_Col-Input-6h-2 | 0.992 |
| Col-Input-24h-1_vs_Col-Input-24h-2 | 0.989 |
| Col-Input-48h-1_vs_Col-Input-48h-2 | 0.912 |
| Col-Input-72h-1_vs_Col-Input-72h-2 | 0.968 |
| Col-H3K27me3-0h-1_vs_Col-H3K27me3-0h-2 | 0.997 |
| Col-H3K27me3-6h-1_vs_Col-H3K27me3-6h-2 | 0.977 |
| Col-H3K27me3-24h-1_vs_Col-H3K27me3-24h-2 | 0.996 |
| Col-H3K27me3-48h-1_vs_Col-H3K27me3-48h-2 | 0.942 |
| Col-H3K27me3-72h-1_vs_Col-H3K27me3-72h-2 | 0.990 |
| *ref6*$^{c}$-Input-0h-1_vs_*ref6*$^{c}$-Input-0h-2 | 0.791 |
| *ref6*$^{c}$-Input-6h-1_vs_*ref6*$^{c}$-Input-6h-2 | 0.898 |
| *ref6*$^{c}$-Input-24h-1_vs_*ref6*$^{c}$-Input-24h-2 | 0.958 |
| *ref6*$^{c}$-Input-72h-1_vs_*ref6*$^{c}$-Input-72h-2 | 0.988 |
| *ref6*$^{c}$-H3K27me3-0h-1_vs_*ref6*$^{c}$-H3K27me3-0h-2 | 0.947 |
| *ref6*$^{c}$-H3K27me3-6h-1_vs_*ref6*$^{c}$-H3K27me3-6h-2 | 0.989 |
| *ref6*$^{c}$-H3K27me3-24h-1_vs_*ref6*$^{c}$-H3K27me3-24h-2 | 0.996 |
| *ref6*$^{c}$-H3K27me3-72h-1_vs_*ref6*$^{c}$-H3K27me3-72h-2 | 0.992 |
| REF6-HA-0h-1_vs_REF6-HA-0h-2 | 0.948 |
| REF6-HA-6h-1_vs_REF6-HA-6h-2 | 0.982 |
| REF6-HA-24h-1_vs_REF6-HA-24h-2 | 0.979 |
| REF6-HA-48h-1_vs_REF6-HA-48h-2 | 0.980 |
| REF6-HA-10d-1_vs_REF6-HA-10d-2 | 0.989 |
